# *In vitro* phenotypic and transcriptomic variation in *Neisseria musculi* morphotypes correlate with colonization variability and persistence *in vivo*

**DOI:** 10.1101/2022.02.03.479073

**Authors:** Eliza Thapa, Leah Lauderback, Cassandra Simmons, Donald L. Holzschu, Adonis D’Mello, Mancheong Ma, Magdalene So, Hervé Tettelin, Nathan J. Weyand

**Affiliations:** Department of Biological Sciences, Ohio University, Athens, Ohio, USA; Department of Microbiology and Immunology, Institute for Genome Sciences, University of Maryland School of Medicine, Baltimore, Maryland, USA; Department of Immunobiology and BIO5 Institute, University of Arizona, Tucson, Arizona, USA; The Infectious and Tropical Disease Institute, Ohio University, Athens, Ohio, USA; Molecular and Cellular Biology Program, Ohio University, Athens, Ohio, USA

## Abstract

Asymptomatic colonization of the upper respiratory tract is a common trait of the two human restricted pathogens, *Neisseria gonorrhoeae* and *Neisseria meningitidis. In vivo* models of pathogenic neisserial infections are heterologous systems that permit short-term colonization but do not fully recapitulate infections in humans. Studying *Neisseria musculi* (Nmus), an oral commensal, in laboratory mice allows investigation of *Neisseria*-host interactions that avoids host restriction barriers. Nmus produces smooth and rough morphotypes on solid media. We compared the *in vitro* phenotypes, biofilm transcriptomes, *in vivo* colonization patterns and burdens of the two Nmus morphotypes. We observed that the two morphotypes differ in biofilm formation, pilin production, transformation frequency, and aggregation *in vitro*. These phenotypes strongly correlated with differential expression of a set of genes in the Nmus biofilms including those that encoded factors for bacterial attachment. *In vivo*, the smooth morphotype stably colonized the oral cavities of all inoculated A/J and C57BL/6J mice at higher burdens relative to the rough. Interestingly, both morphotypes colonized the oral cavities of A/Js at higher magnitudes than in C57BL/6Js. Gut colonization by the smooth morphotype was qualitatively higher than the rough. Nasal colonization in the A/Js were transient following nasal inoculations. Collectively, our results demonstrate that colonization by Nmus can be affected by various factors including Nmus morphotypes, inoculation routes, anatomical niches, and host backgrounds. The Nmus-mouse model can use variable morphotype-host combinations to study the dynamics of neisserial asymptomatic colonization and persistence in multiple extragenital niches.

**IMPORTANCE:** Animal models for human adapted pathogenic *Neisseria* spp. do not fully mimic human infections and are complicated by host restriction barriers that can hinder long-term persistence. Such barriers can be avoided by studying *Neisseria* spp. native to the animal host used for disease models. *Neisseria musculi* (Nmus) isolated from wild mice colonizes the oral cavity and gut of laboratory mice for extended periods. Nmus shares host interaction factors with species pathogenic to humans and thus provides a native system to study orthologs of factors that may facilitate asymptomatic colonization and persistence in the human upper respiratory tract. We investigated the Nmus-mouse system to compare *in vitro* and *in vivo* phenotypes of two Nmus morphotypes. Our results support the hypothesis that the two morphotypes vary in different aspects of *Neisseria*-host interactions. Future use of the Nmus-host system will help identify molecular mechanisms required for neisserial asymptomatic colonization, dissemination, and persistence.

## INTRODUCTION

At least 30 species of *Neisseria* inhabit a wide range of mammalian and non-mammalian hosts (1). Most of these species are professional colonizers of mucosal surfaces where they reside as commensals. In humans, *Neisseria* species are well adapted commensals of the oral and nasopharyngeal niches and are believed to be early colonizers of these sites (2-4). The two best studied *Neisseria* species - *Neisseria gonorrhoeae* and *Neisseria meningitidis* (Nme) cause severe infections only in humans (5). While *N. gonorrhoeae* is an obligate pathogen that causes gonorrhea, Nme is called an “obligate commensal” or “accidental pathogen” (6). Both pathogens can asymptomatically colonize multiple anatomical niches in humans (7, 8).

Over 12 commensal *Neisseria* spp. are components of the human microbiota, however they are rarely studied (9). Human commensal *Neisseria* spp. are known to impact the survival and presence of their disease-causing counterparts. A study conducted using human volunteers show that *Neisseria lactamica*, a nasal commensal can displace Nme and hinder its acquisition (10). Another commensal, *Neisseria cinerea* is known to affect the motility, microcolony formation and epithelial cell attachment of Nme (11). Furthermore, using a vaginal mouse model of infection, Kim et. al., showed that commensal *Neisseria* can kill human *Neisseria* pathogens by a DNA-dependent mechanism (12). Commensal *Neisseria* spp. share a collection of genes with pathogens (13). Since commensal and pathogenic *Neisseria* spp. inhabit common niches, studying commensals will help determine if they share similar persistence mechanisms with the pathogens.

*Neisseria musculi* (Nmus) is a commensal of the oral cavity of wild caught mice. It can easily be grown *in vitro* and encodes orthologs of several host-interaction factors found in human pathogenic *Neisseria* spp. (14). The Nmus-mouse model lacks host-restriction barriers that hinder many human pathogenic *Neisseria* infection models and will allow investigation of the principles of neisserial asymptomatic carriage and persistence. Nmus produces two morphotypes when grown on solid media - smooth and rough. Using the Nmus rough morphotype, Ma et.al., showed that Nmus can colonize the oral cavity and gut of laboratory mice for up to a year (15). Colonization was natural without the use of any antibiotics or hormones and the animals remained healthy throughout the study. The susceptibility to Nmus colonization is affected by host genetics and the innate immune system (15, 16). Furthermore, colonization by Nmus induces components of adaptative immune responses that influence host colonization susceptibility (16).

In this study, we further explored the Nmus-mouse system by comparing the *in vitro* phenotypes and colonization abilities of the smooth and rough Nmus morphotypes. Using *in vitro* experiments and transcriptomic analysis, we show that the two morphotypes differ in biofilm formation, pilin production, aggregation, vancomycin susceptibility and expression of a repertoire of genes including those involved in Type IV pilus (Tfp) biogenesis. The Tfp biogenesis loci in Nmus shared local synteny with Nme strain 8013. Furthermore, we hypothesized that Tfp related *in vitro* phenotypic differences influence *in vivo* persistence phenotypes for both morphotypes. We observed that the smooth morphotype can efficiently and persistently colonize the oral cavity of every inoculated mouse at a higher burden than the rough in two mouse strains. Higher oral burdens of both Nmus morphotypes were observed in A/J mice compared to C57BL/6J mice. Similarly, in the gut, a higher frequency of mice was colonized by the smooth morphotype. Only a transient colonization was noted in the nasal cavity. Moreover, Nmus morphotypes can disseminate from the nasal cavity to oral niches followed by stable oral colonization. Nmus morphotypes, anatomical niches and host backgrounds can all influence *in vivo* persistence phenotypes. Thus, morphotype-host combinations can be used to focus research toward a range of queries about *Neisseria*-host interactions.

## RESULTS

### Nmus morphotypes form disparate biofilms

Nmus morphotypes form biofilms on glass (14). To further study the nature of biofilms on abiotic surfaces, static biofilms on polystyrene were assayed using Crystal Violet staining (17) with some modifications as described in the materials and methods. We observed that the smooth morphotypes formed nearly 2-fold less biomass *(P* < 0.01) compared to the rough (Fig. 1B). Interestingly, we also noted that smooth biofilms were more adhesive than rough biofilms which were very fragile (Fig. 1C). Biofilm fragility assays showed that the smooth biofilms were more resistant to saline washes than the rough (*P* < 0.001) (Fig. 1D). This suggests that the smooth morphotype forms thinner biofilms which are sticky and attach more firmly to plastic surfaces than fragile rough biofilms.

**FIG 1.**
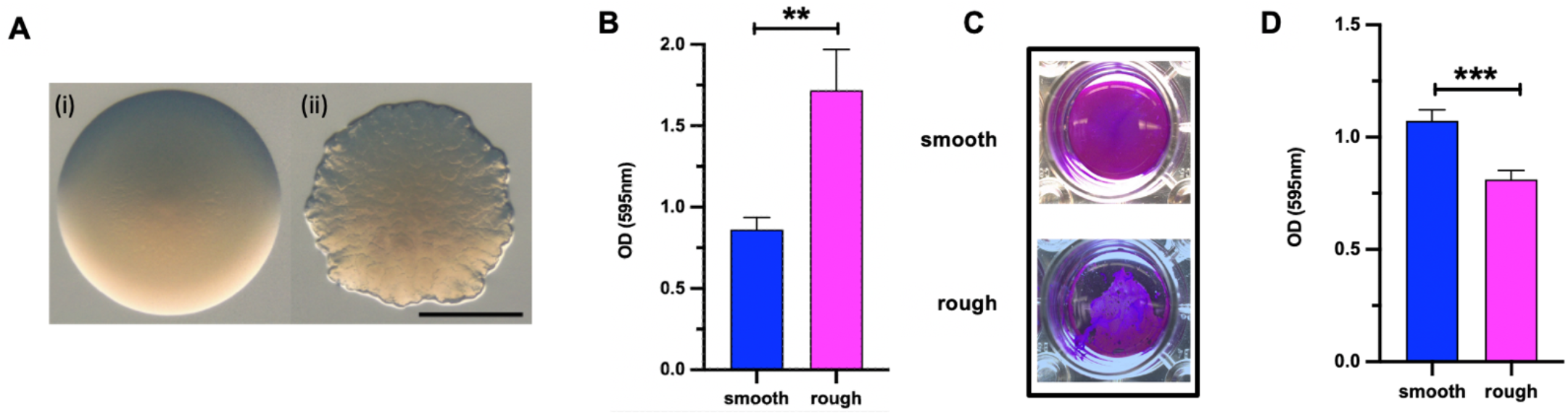
Nmus morphotype biofilms. (A) Smooth (i) and rough (ii) Nmus colony morphotypes on GCB agar after 48 h. Scale bar: 0.5 mm. (B) Quantification of Nmus biofilm formation. Biofilm fragility assay images (C) and quantitation (D). Experiments were performed three times with technical triplicates (B) or quadruplicates (C, D). Error bars indicate SEM. Statistical analysis was determined by the Student’s *t* test. **, *P* < 0.01; ***, *P* < 0.001.

### Transcriptome sequencing of Nmus morphotype biofilms

We hypothesized that the phenotypic differences observed in morphotype biofilms may be due to altered gene expression. To test this, we performed transcriptome analysis (RNA-Seq) on RNA isolated from static morphotype biofilms. Principal Component Analysis (PCA) was used to interrogate whole transcriptome profile similarities across biological replicates. For both Nmus morphotypes, only two of the three replicates clustered together (Fig. 2A). Outliers were excluded for differential expression (DE) calculations between Nmus biofilms (Fig. 2B and Data Set S1). Of note, DE gene results using all three replicates showed similar trends for the most biologically relevant genes further scrutinized in this study (Fig. S1 and Data Set S2).

**FIG 2.**
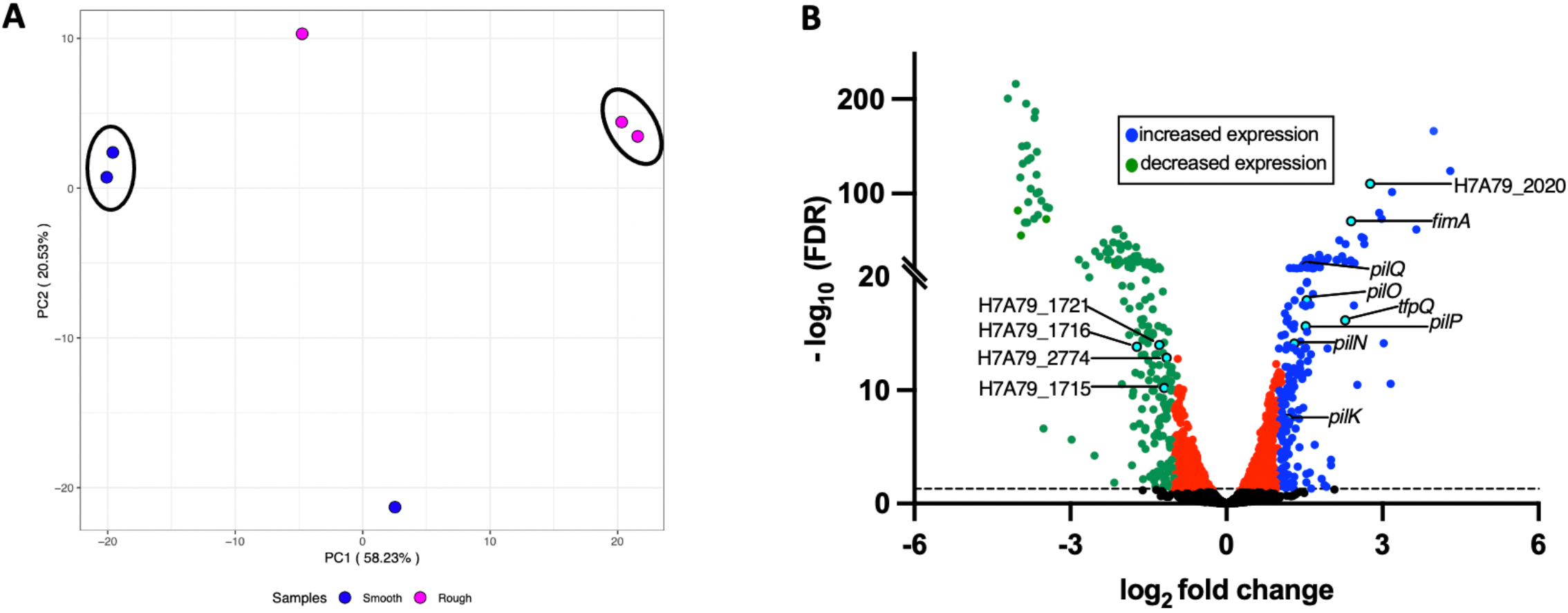
RNA-Seq analysis of Nmus morphotype biofilms. (A) Principal Component Analysis of Nmus biofilm gene expression profiles. Biological replicates used for RNA-Seq differential gene expression analysis are outlined by ovals. (B) Volcano plot of differentially expressed genes in Nmus biofilms. Dotted line indicates a False Discovery Rate (FDR) threshold of 0.05. Blue and green dots are genes with significant FDR and log_2_ fold changes for increased (> 1) and decreased (< -1) expression in smooth biofilms, respectively. Black dots and red dots represent genes that do not reach the FDR and log_2_ fold change thresholds, respectively. Teal dots with annotations on the right indicate genes that encode factors associated with attachment including Tfp. Teal dots with annotations on the left denote locus tags for genes that encode putative efflux pumps.

Using two replicates for each morphotype resulted in 413 genes that were differentially expressed between the two biofilms with a log_2_ fold-change < = −1 or > = 1 and significant False Discovery Rate (FDR) < = 0.05. Among the DE genes, 201 genes had higher expression in the smooth biofilm compared to the rough biofilm (Fig. 2B). Interestingly, eight genes more highly expressed in the smooth biofilm encoded factors that can influence attachment of *Neisseria* spp. or other bacterial species to biotic and abiotic surfaces (Table 1). These included fimbrial and hemagglutinin family proteins and Type IV pilus (Tfp) biogenesis proteins. Tfp are multi-subunit nanomachines involved in attachment, colonization, transformation, and motility in *Neisseria* spp. (18). Of the Tfp biogenesis genes, ten were more highly expressed in smooth biofilm (5 passing both significance cutoffs), including *pilX*, a minor pilin (Tables 1 and S1). These gene expression differences correlate with published adherence functions for their encoded proteins (19, 20) and with our observations that the smooth biofilms are more strongly attached to polystyrene surfaces than rough biofilms.

**TABLE 1.** Differentially expressed genes associated with adhesion in Nmus biofilms

| Gene ID/Locus tag | Accession no. in Nmus | Protein ID | log<sub>2</sub>fold change<sup>c</sup> | -log (FDR) |
| --- | --- | --- | --- | --- |
| *fimA*<sup>a</sup> | QNT58552.1 | fimbrial protein | 2.40 | 70.65 |
| H7A79_2020<sup>a</sup> | QNT59712.1 | hemagglutinin family protein | 2.76 | 110.35 |
| *pilQ*<sup>b</sup> | QNT57933.1 | type IV pilus biogenesis protein PilQ | 1.54 | 23.05 |
| *pilP*<sup>b</sup> | QNT58312.1 | type IV pilus biogenesis protein PilP | 1.52 | 15.64 |
| *pilO*<sup>b</sup> | QNT58822.1 | type IV pilus biogenesis protein PilO | 1.54 | 17.95 |
| *pilN*<sup>b</sup> | QNT59588.1 | type IV pilus biogenesis protein PilN | 1.30 | 14.10 |
| *tfpQ*<sup>a</sup> | QNT59545.1 | fimbrial protein Q | 2.28 | 16.18 |
| *pilK*<sup>b</sup> | QNT59870.1 | type IV pilus biogenesis protein PilK | 1.19 | 7.49 |

<sup>a</sup> Gene ID/ locus tag and protein ID are retrieved from the Nmus genome (GenBank: CP060414.1)
<sup>b</sup> Gene and protein ID are retrieved based on homology to Tfp biogenesis factors encoded by the Nme genome (Genbank: FM999788.1)
log<sub>2</sub>foldchange indicates increased gene expression in the smooth Nmus biofilms compared to the rough biofilms

### Type IV pilus biogenesis operons of Nmus and Nme strain 8013 have shared synteny

To identify orthologous clusters of Tfp biogenesis genes in Nmus, we used the Pan-genome ortholog clustering tool (PanOCT) (21). Since gene annotations and functional characterization of Tfp biogenesis genes have been extensively performed in Nme strain 8013, its genome was used for the analysis (22, 23). We observed local synteny of Tfp biogenesis loci between Nmus and Nme (Fig. 3). Some differences included: (i) a gene encoding a helix-turn-helix domain protein located between *ispG* and *pilW* (labeled ORF1) was absent in Nme, (ii) Nmus *comP* and *pilV* genes were located together but found at separate loci in Nme, and (iii) a gene encoding a YGGT family protein located between *proC* and *dksA* (labeled ORF2) was absent in Nme. Additionally, analysis of proteins encoded by syntenic genes using reciprocal BLASTp showed query coverage of at least 80% and identity between 37-90% (Table S2).

**FIG 3.**
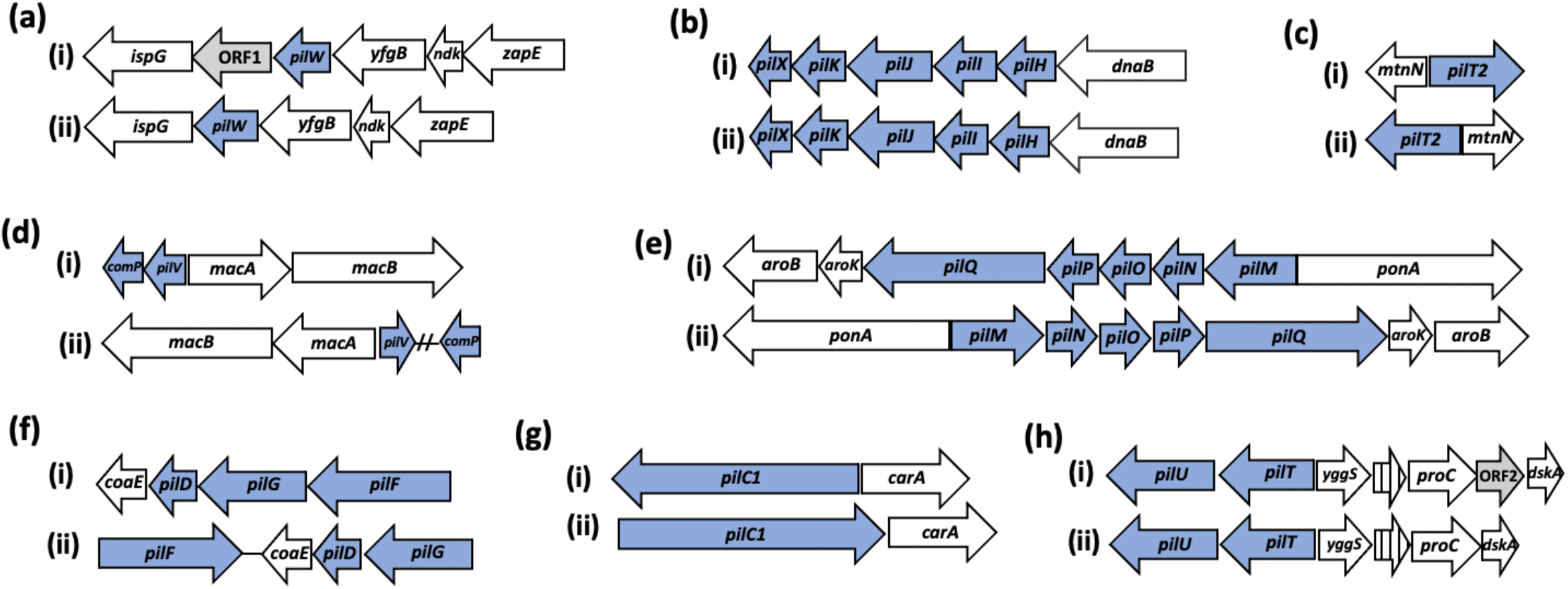
Tfp biogenesis gene operons in Nmus and Nme 8013 show conserved synteny. Comparison of Type IV pilus biogenesis gene loci (a-h) between (i) Nmus strain 831 and (ii) Nme strain 8013 (see also Table S2). Type IV pilus biogenesis genes are indicated by blue arrows and flanking genes sharing synteny by white arrows; grey arrows represent strain-specific genes. Gene symbols for syntenic genes were retrieved from previously annotated Nme genomes (see methods). Hatched lines in panel (d) indicate a region of greater than 300 kb. Stripped white arrows in (h) encode putative RNA-binding proteins.

Interestingly, our PanOCT analysis did not detect synteny for *pilE*, which encodes the major Tfp pilin in Nme. We used an alternative approach, the PilFind program (24), to identify potential pilins in the Nme and Nmus genomes. PilFind identified eight putative pilins in Nme and seven in Nmus including two encoded by *tfpQ* (H7A79_1926) and *fimA* (H7A79_1928) (Table S3). Since higher expression of *fimA* and *tfpQ* were detected in the smooth morphotype, we hypothesize either could encode a major pilin. Furthermore, multiple sequence alignments and phylogenetic analysis of pilin proteins found that the major Nme 8013 pilin, PilE, was more closely related to FimA and TfpQ of Nmus than the minor pilins PilX, ComP or PilV (Fig S2.). Future experiments will be needed to confirm if either gene encodes the major pilin expressed in Nmus morphotypes.

In conclusion, the highly conserved synteny and protein identity of Tfp biogenesis loci in Nmus and Nme suggests that Nmus will serve as a good model organism for the study of *in vivo* functions of Tfp.

### Smooth morphotype gene expression correlates with Tfp-dependent functions

Since several Tfp biogenesis genes were differentially expressed in Nmus morphotypes (Fig. 2B and Table S1), we evaluated *in vitro* phenotypes that can correlate with Tfp function. Pilin from Nmus strains was extracted by shearing and ultracentrifugation. We detected a band from the smooth morphotype preparations with a molecular mass of ∼16 kDa (Fig. 4A). This mass is similar to the major pilin in *Neisseria* spp. (25). While a visible band with similar mass was not detected from rough morphotype crude pilin preparations, the rough morphotype does express both putative pilin genes *fimA* and *tfpQ*, albeit at lower levels than the smooth morphotype (Table 1). As controls we used *pilT* deletion strains in both morphotype backgrounds. Since, Δ*pilT* strains cannot retract their pili, they typically yield more pilin (26). We observed that both Δ*pilT* Nmus strains had more intense pilin bands compared to the wild type strains. Further analyses will be required to identify which genes encode the pilins detected in Fig. 4A.

**FIG 4.**
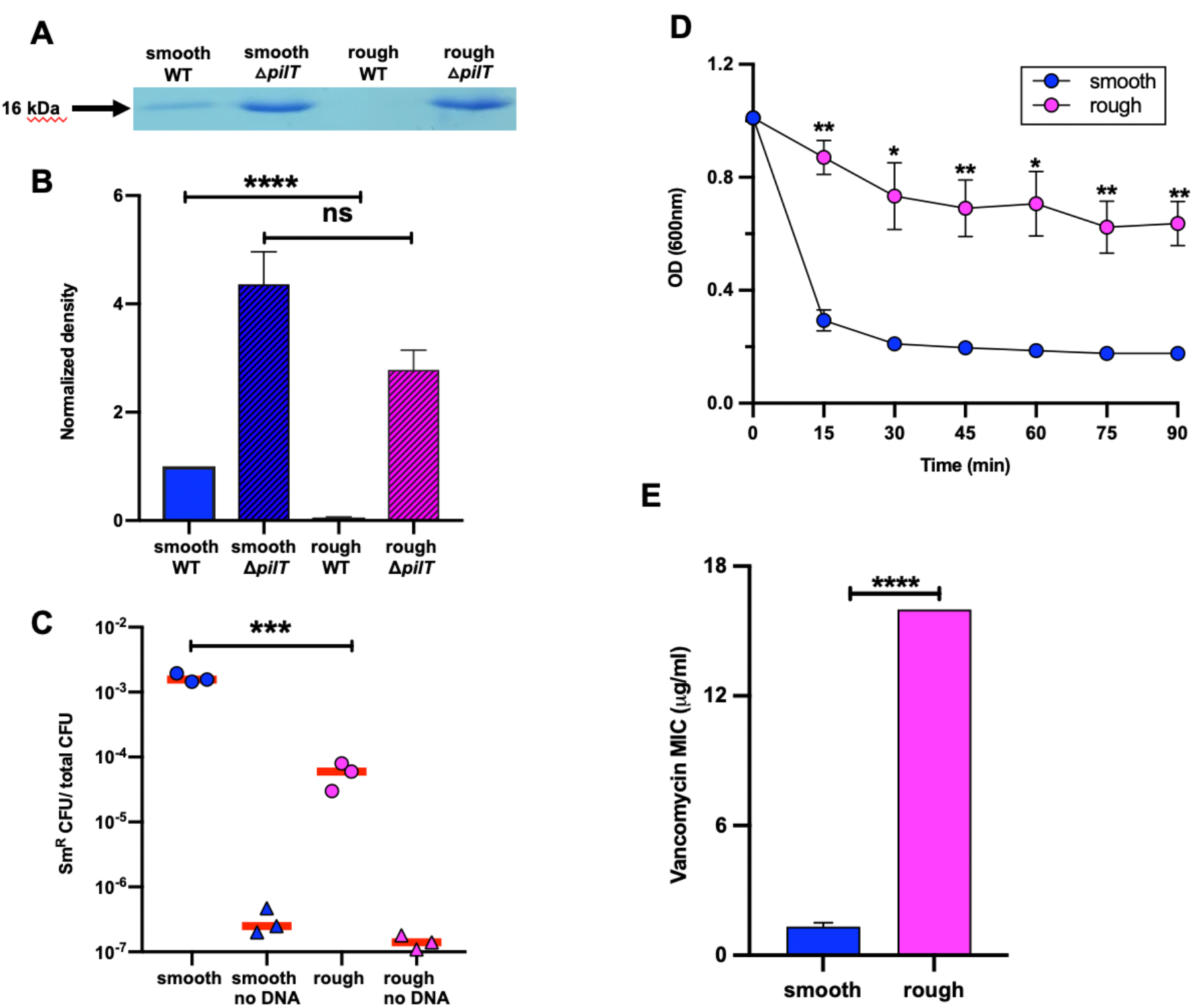
*In* vitro phenotypes of Nmus morphotypes. (A) Commassie blue staining of pilin extracted from Nmus morphotypes. (B) Quantitation of pilin produced by Nmus strains. Nmus Δ*pilT* strains were used as controls. WT, wild type. (C) Transformation frequencies of Nmus morphotypes. Assays without added donor DNA were used as negative controls. (D) Aggregation of Nmus morphotypes. (E) Vancomycin MICs of Nmus morphotypes. Red lines indicate means. Error bars indicate SEM from three independent experiments (B-E). Statistical analyses for all experiments were performed using the Student’s *t* test. *,*P* < 0.05; **, *P* < 0.01; ***, *P* < 0.001; ****, *P* < 0.0001; ns, *P* > 0.05.

Since Tfp plays a major role in the natural competence of *Neisseria* spp. (27, 28), we measured the transformation frequencies of both morphotypes. Genomic DNA from AP2098, a streptomycin resistant (Sm^R^) Nmus strain was used as donor DNA while assays without added donor DNA served as negative controls (15). The transformation frequency of the smooth morphotype was ∼30 times higher than the rough morphotype In the absence of DNA, the frequencies of the spontaneous Sm^R^ smooth and rough morphotypes were > 3 logs and >2 logs lower respectively (Fig. 4C).

Helaine et. al demonstrated a role for *pilX* in bacterial aggregation and formation of Nme microcolonies (29). *pilX* of Nmus shares 39% identity with that of Nme strain 8013. The *pilX* gene was expressed 1.8-fold (log_2_ fold change of 0.85) higher in the smooth biofilm compared to the rough (Table S1). We conducted aggregation assays by suspending bacterial lawns in PBS and monitoring the absorbance every 15 min. We observed that within 15 min of the assay, and for up to 90 min, the smooth morphotype showed a significant decrease in aggregation relative to the rough morphotype (*P* < 0.05) (Fig. 4D).

Together, our results indicate that the two morphotypes show significant *in vitro* phenotypic differences that include pilin production, transformation frequencies and aggregation. These *in vitro* phenotypes mimic previously published Tfp-dependent functions and correlate with our differential gene expression data.

### Smooth morphotype gene expression correlates with vancomycin susceptibility

In the smooth biofilms, 212 genes showed less expression by ≥ 2-fold compared to the rough biofilm. Among them, four genes encode different outer membrane efflux pumps (Fig. 2B). One of these genes, H7A79_1716 in the Nmus genome (30), encodes a putative component of the hemolysin translocation system (HTS). *Escherichia coli* expressing a fully functional HTS is susceptible to vancomycin (31). To correlate gene expression differences with vancomycin susceptibility, we measured the vancomycin MICs for the two Nmus morphotypes. We observed that the smooth morphotype had a 16-fold lower vancomycin MIC (1 mg ml^-1^) compared to the rough morphotype (16 mg ml^-1^) (Fig. 4E). Our MIC results provide an intriguing correlation that may indicate that H7A79_1716 or other efflux pump gene expression differences could be responsible for the vancomycin MICs of the Nmus morphotypes.

### Smooth morphotype efficiently colonizes the oral cavity of inoculated mice

Based on our *in vitro* characterization of the two morphotypes, we speculated they may show colonization differences *in vivo*. A previous study by Ma et. al demonstrated that the rough morphotype colonizes the oral cavity and gut of laboratory mice for extended periods (15). Since colonization abilities of the smooth morphotype have not been investigated, we compared the two Nmus morphotypes following oral cavity inoculations of A/J and C57BL/6J (B6) mice.

We observed that following oral inoculations, 100% of A/J mice (16/16) inoculated with the smooth morphotype were persistently colonized in the oral cavity for 10 weeks. In contrast, only 69% (11/16) of mice were colonized with the rough morphotype (*P* < 0.01) (Fig. 5A). Furthermore, we also noted that the smooth morphotype colonized the oral cavity of A/J mice at a higher CFU burden than the rough morphotype (Fig. 5B to D). A significant difference in the mean Areas Under the Curve (mean AUCs) (log_10_ CFU versus time) between the two morphotypes was also observed (*P* < 0.01) (Fig. 5E).

**FIG 5.**
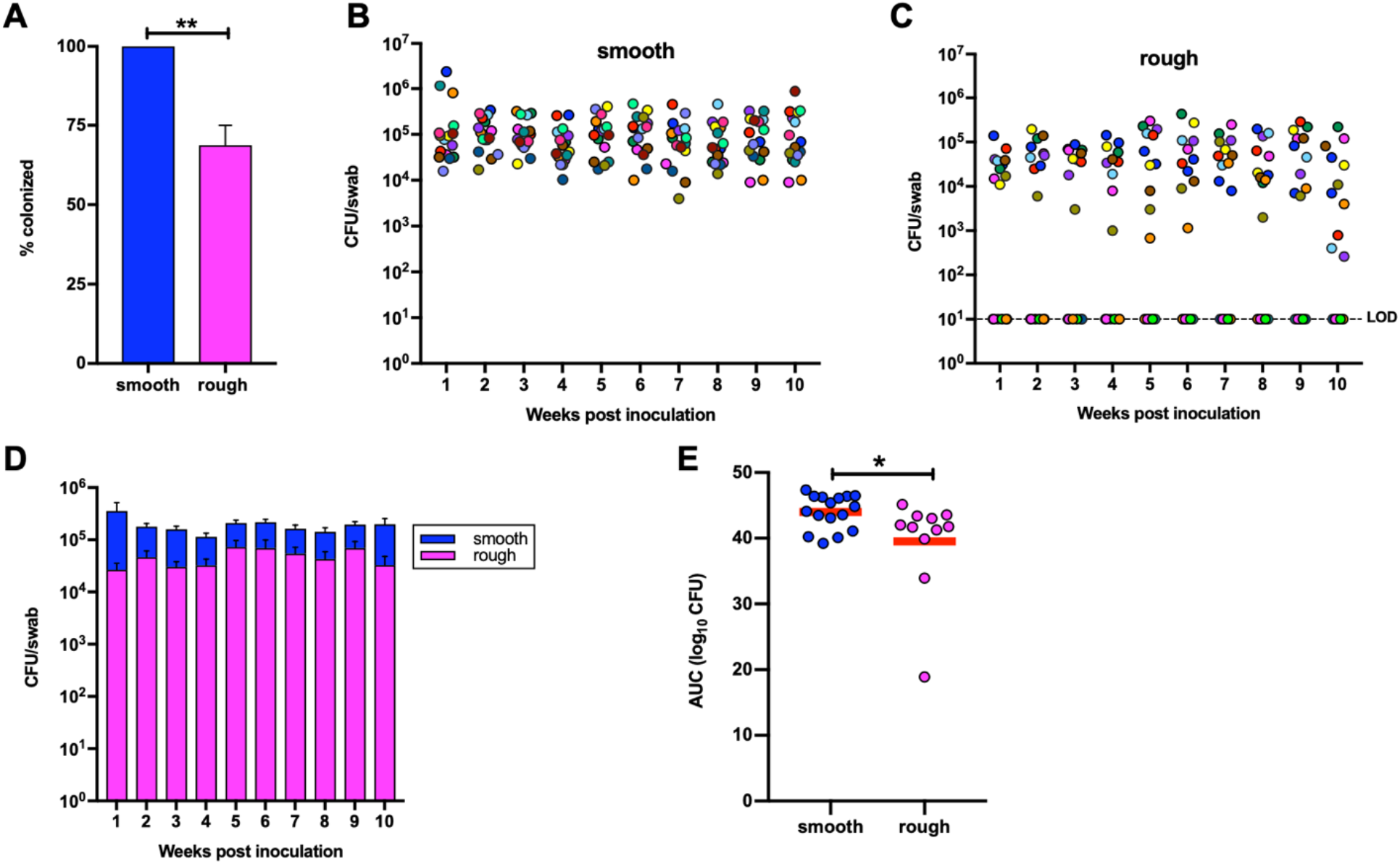
Nmus oral colonization phenotypes in A/J mice. (A) Percentage of A/J mice (n= 16 for both) colonized by Nmus morphotypes following oral inoculations. (B, C) Oral burdens of Nmus morphotypes following oral inoculations. Each colored dot represents a single mouse. LOD, limit of detection. (D) Comparison of the oral burdens of Nmus morphotypes. (E) Cumulative CFU as area under the curve (AUC). The mean AUC (log_10_ CFU versus time) was calculated for each mouse to estimate the bacterial burden over time. The means under the curves were compared between the two morphotypes. Error bars indicate SEM. Red lines indicate means. Statistical analyses for A and D were performed using the Student’s *t* test. *, *P* < 0.05; **, *P* < 0.01.

Similar results were obtained in the B6 mice. When inoculated orally, 100% of B6 mice (12/12) inoculated with the smooth morphotype were colonized relative to 33% mice (4/12) colonized by the rough morphotype (*P* < 0.01) (Fig. S3A). The oral burden of the smooth morphotype was nearly 4-fold higher than the rough. The mean AUCs between the two morphotypes was observed to be significantly different (*P* < 0.01) (Fig. S3).

After 10 weeks of oral colonization, A/J and B6 mice were euthanized, and their tongues harvested to enumerate associated CFU per gram of tissue. Consistent with the observations from oral swabs, tongue CFUs were recovered from 100% mice colonized with the smooth morphotype. Similarly, only those mice with detectable oral colonization with the rough morphotype (69% A/Js and 33% B6s) showed tongue CFUs (Fig. S4). The tongue CFU burdens were qualitatively higher for the smooth morphotype compared to the rough in both mouse backgrounds.

To test if the host background affects the colonization ability of Nmus, we compared the oral colonization burdens of Nmus morphotypes between A/J and B6 mice. Our results showed that both morphotypes colonize the oral cavities of A/J mice at higher burdens than in B6 mice (Fig. 6A and C). Significant differences in the mean AUCs were observed for both Nmus morphotypes between A/J and B6 mice (Fig. 6B and D).

**FIG 6.**
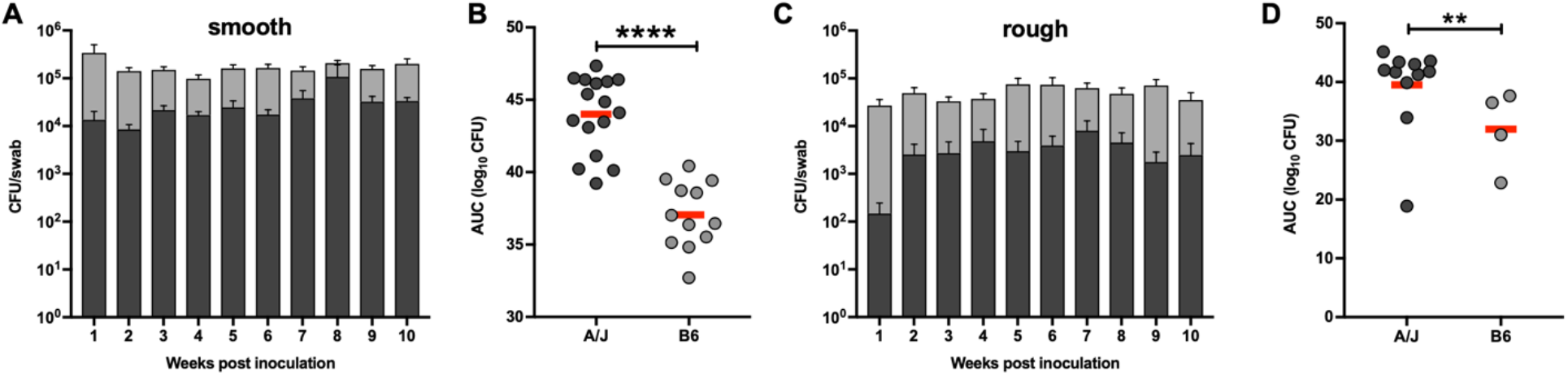
Comparison of Nmus morphotypes oral burdens between A/J and B6 mice. Oral burdens of Nmus morphotypes in A/J (A) and B6 (C). Cumulative CFU as area under the curve (AUC) for smooth (B) and rough (D) Nmus morphotypes. The mean AUC (log_10_ CFU versus time) was calculated for each mouse to estimate the bacterial burden over time. The means under the curves were compared between the two morphotypes. Error bars indicate SEM. Red lines indicate means. Statistical analyses for B and D were performed using the Student’s *t* test. **, *P* < 0.01; ****, P < 0.0001.

In summary, these results suggest that the smooth morphotype more consistently colonizes the oral cavity of inoculated mice and at higher burdens than the rough. Furthermore, both morphotypes can orally colonize the A/J mice at higher CFU burdens than B6 mice.

### Smooth morphotype colonizes the guts of mice at higher frequency

Following oral inoculations, we monitored the persistence of the Nmus morphotypes in the guts of A/J and B6 mice. Notably, the frequency of mice colonized in the gut was lower compared to the oral colonization in both mouse backgrounds. Nevertheless, like in the oral cavity, both mouse strains were more stably colonized with the smooth morphotype than with the rough. A higher percentage of A/J mice were colonized in the gut by the smooth morphotype than the rough (Fig. 7A). A similar trend was seen when gut colonization was monitored in B6 mice (Fig. S5A). Furthermore, a qualitative difference in the mean AUC was observed between the two morphotypes in both mouse backgrounds (Fig. 7D and S5D). This implies that although the gut colonization by Nmus morphotypes is less frequent than in the oral cavity, the smooth morphotype can colonize this niche at a higher frequency than the rough.

**FIG 7.**
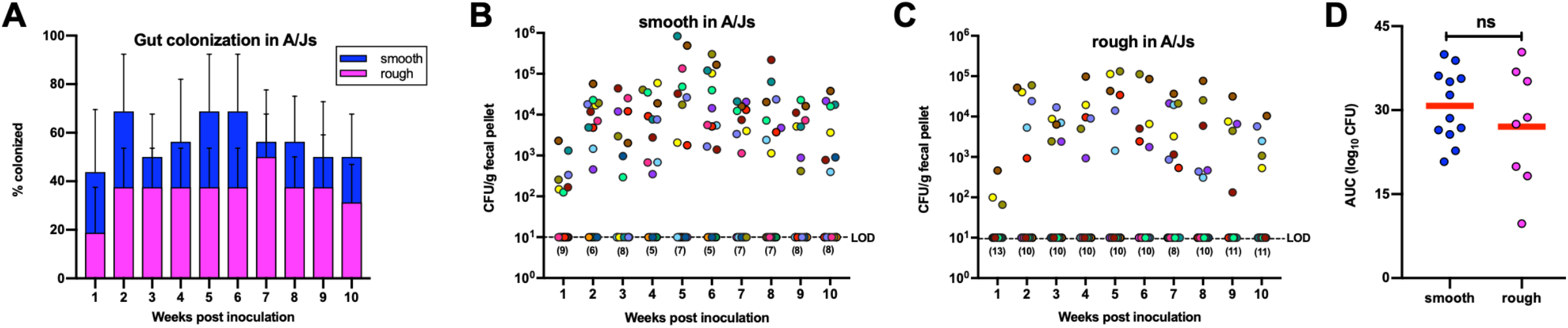
Gut colonization of Nmus morphotypes in A/J mice following oral inoculations. (A) Percentage of A/Js colonized by Nmus morphotypes in the gut. Data represent mean with SEM from four independent experiments. Fecal burdens of smooth (B) and rough (C) morphotypes in A/Js (n= 16 for both). Each colored dot represents a single mouse. Numbers in parentheses represent the number of mice in which colonization was not detected. LOD, limit of detection. (D) Cumulative CFU as area under the curve (AUC). The mean AUC (log_10_ CFU versus time) was calculated for each mouse to estimate bacterial burden over time. Red lines indicate means. Statistical analysis for D was performed using the Student’s *t* test. ns, *P* > 0.05.

### Nmus transiently colonizes the nasal cavity of A/J mice

Next, we investigated the ability of Nmus morphotypes to colonize the nasal cavities of A/J mice. Unlike the oral cavities, the smooth morphotype did not colonize the nasal cavities of 100% of the inoculated A/Js. However, notable differences in colonization and burdens between the two morphotypes were observed in the nasal cavities. This observation was true for both time points analyzed. At 4 dpi, a significant difference in nasal colonization was observed between the two morphotypes (67% vs 0%). The rough morphotype was not detected at 4 dpi. At 8 dpi, only a qualitative difference in nasal colonization was observed (57% vs 14%) (Fig. 8A).

**FIG 8.**
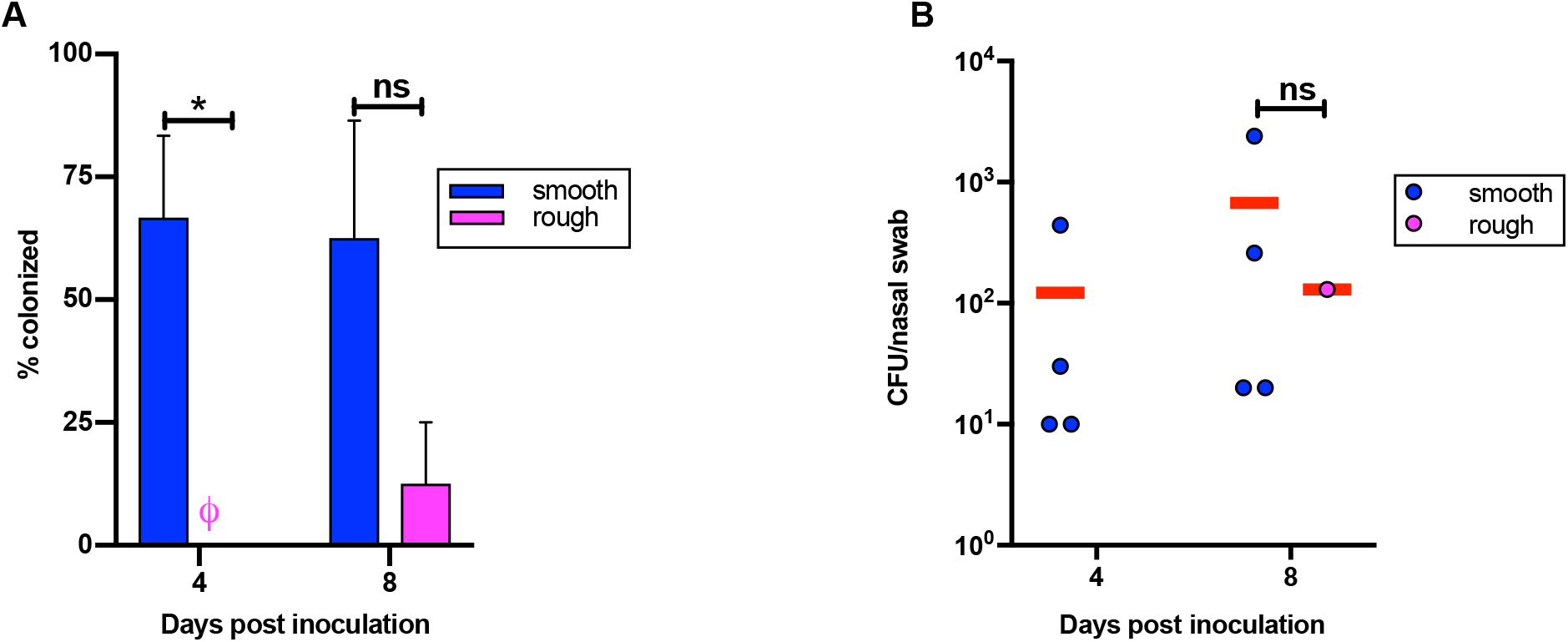
Nmus nasal colonization phenotypes in A/J mice. (A) Percentage of A/J mice colonized in the nasal cavity by Nmus morphotypes after 4 days post inoculation (dpi) and 8 dpi. Data represents mean with SEM. ø, not detected. (B) Nasal burdens of Nmus morphotypes following 4 dpi and 8 dpi. Red lines indicate mean bacterial burdens in the nasally colonized mice. Experiments were performed three or four times with n=6 mice for 4 dpi and n=7 for 8 dpi. Statistical analyses were performed using the Student’s *t* test for (A) at 4 dpi and Mann-Whitney’s U-test at 8 dpi for (A) and (B). *, *P* < 0.05; ns, *P* > 0.05.

We also observed differences in bacterial burdens between the nasal and oral cavities in A/J mice. The average nasal burdens of the smooth morphotype in the colonized A/Js were ∼100 fold lower than the average oral burdens (Fig. 5B and 8B). Similarly, the average nasal burdens for the rough morphotype were ∼1000-fold lower than the average oral burdens (Fig. 5C and 8B). Furthermore, in the nasal cavities, the average smooth burdens were qualitatively higher than rough at 8 dpi (Fig. 8B). Our attempts to study the nasal colonization for extended periods were unsuccessful as we did not detect Nmus in nasal swabs collected at 15 or 30 dpi (data not shown). Thus, we conclude that Nmus morphotypes have limited ability to colonize the nasal cavities of A/Js and the smooth morphotype can colonize the nasal cavities of a higher percentage of inoculated mice compared to the rough.

### Nmus can disseminate to the oral cavity and gut of A/J mice following nasal inoculations

We further investigated the ability of Nmus to disseminate to the oral cavities and guts of A/J mice following nasal inoculations. We noted a significant difference in oral colonization by the two morphotypes following nasal inoculations. While 100% mice inoculated with the smooth morphotype were orally colonized at 3 dpi (12/12) and 7 dpi (10/10), only 33% (4/12) and 40% (4/10) mice inoculated with rough were colonized at 3 dpi (*P* < 0.01) and 7 dpi (*P* < 0.05) respectively (Fig. 9A). We also observed a significant difference in the oral burdens between the smooth and rough morphotypes in nasally inoculated mice. After 3 dpi, the smooth morphotype colonized the oral cavity ∼ 8-fold higher than the rough (*P* < 0.01). Similarly, at 7 dpi, the average oral burdens of smooth morphotype were ∼33 fold higher than the rough (*P* <0.0001) (Fig. 9B). A significant difference between mean AUCs (log_10_ CFU versus time) was also observed between the two morphotypes at both 3 dpi and 7 dpi (Fig. 9C). Furthermore, the tongue CFUs of the smooth morphotype were qualitatively higher than the rough (Fig. S6A) following nasal inoculations.

**FIG 9.**
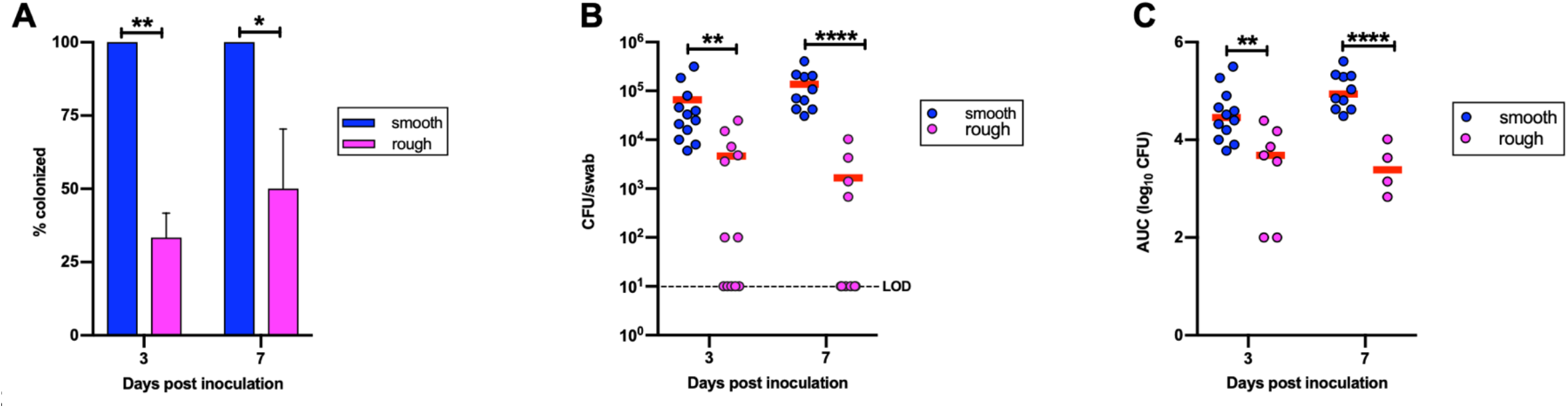
Nmus oral colonization phenotypes in A/J mice following nasal inoculations. (A) Percentage of A/J mice orally colonized after 3 days post inoculation (dpi) and 7 dpi. Data represents mean with SEM. (B) Oral burdens of Nmus morphotypes following nasal inoculations at 3 dpi and 7 dpi. LOD, limit of detection. (C) Cumulative CFU as area under the curve (AUC). The mean AUC (log_10_ CFU versus time) was calculated for each colonized mouse to estimate the bacterial burdens at 3 dpi and 7 dpi. The means under the curves were compared between morphotypes. Experiments were performed three or four times with n=12 mice for 3 dpi and n=10 for 7 dpi, respectively. Each dot in (B and C) represent bacterial counts obtained from a single mouse. Red lines indicate means. Statistical analyses were performed using the Student’s *t* test for (A) and (C), and on log transformed data for (B). *, *P* < 0.05; **, *P* < 0.01; ****, *P* < 0.0001.

Gut colonization of nasally inoculated mice mimicked the trend following oral inoculations. Namely, more mice inoculated with the smooth morphotype were colonized in the gut compared to the rough. The fecal burdens for the smooth morphotype were only qualitatively higher than the rough (Fig. S6B). Taken together, these results indicate that both Nmus morphotypes can disseminate to the intestinal niche irrespective of oral or nasal inoculations.

## DISCUSSION

In this study, we have presented several *in vitro* and in *in vivo* differences between the smooth and rough morphotypes of Nmus. Using transcriptomic data, we have identified a subset of Tfp biogenesis genes and efflux pumps that are differentially expressed in the biofilms of the two morphotypes. Several *in vitro* phenotypes influenced by Tfp, and antibiotic susceptibility positively correlated with the transcriptomic data. We observed significant *in vivo* differences in the colonization abilities and burdens of Nmus morphotypes in the oral cavities, nasal cavities and guts of laboratory mice. Nmus persistence phenotypes vary in different anatomical niches and are influenced by the mouse background used, suggesting that this model system can be used to study various aspects of neisserial colonization.

Phenotypic variation is an important adaptive mechanism for some bacterial pathogens for successful colonization and infection (32). Persistence phenotypes can strongly correlate with differences in colony morphologies and associated genetic and physiological traits (33). Colony morphotypes of *Haemophilus influenzae* (34) and *Porphyromonas gingivalis* (35) have differential colonization abilities and antibiotic resistance profiles, respectively. In the case of Nmus, although the mechanisms contributing to the different colony morphologies are still being investigated, it may be dependent upon phase variation of specific factors as observed in other *Neisseria spp*. (36, 37).

Our RNA-Seq analysis identified differential expression of several genes between the biofilms produced by the Nmus morphotypes. Five genes that significantly showed higher expression in the smooth biofilms encode proteins involved in Tfp biogenesis (Table 1). These genes are *pilK, pilN, pilO, pilP* and *pilQ*. In Nme, at least 23 proteins are involved in pilus biogenesis and function (38). PilK, PilN, PilO and PilP are structural proteins required for pilus assembly. PilQ is an outer membrane secretin involved in translocation of pilin proteins and can mediate DNA uptake during transformation (39, 40). Our *in vitro* results for the smooth morphotype correlate well with the differential gene expression data and known functions of Tfp proteins in pilin production and transformation. Genes encoding pilus proteins, PilX and PilC1 were also differentially expressed in the two biofilms although they did not meet our cut off criteria for fold change (Table S1). PilX, a minor pilin in *Neisseria* spp. was expressed more in the smooth biofilm by log_2_ fold change of 0.85 compared to the rough. PilX confers aggregative properties in Nme (29). PilC1 also showed more expression in the smooth biofilm by log_2_fold change of 0.98 and a significant FDR. In human pathogenic *Neisseria* spp., PilC1 facilitates bacterial adhesion to the host cells (41-43). The increased expression of these genes in the smooth morphotype correlates with its rapid aggregation and increased biofilm strength and attachment on the abiotic surfaces respectively. Although, a direct role of Tfp associated genes in Nmus was not demonstrated, our study has helped to develop testable hypotheses that will lead to a better understanding of the function of Tfp proteins *in vivo*.

Efflux pumps frequently function to confer antibiotic resistance in Gram-negative bacteria (44). Among the four efflux pump genes that have lower expression in the smooth biofilms, orthologs of the hemolysin translocation protein are associated with vancomycin resistance (31, 45). *In vitro*, the lower expression of this efflux pump in the smooth biofilms correlated with significantly lower vancomycin MIC of this morphotype. Although other resistance mechanisms may exist, our results strongly suggest that the vancomycin resistance profile in Nmus morphotypes may be related to differential expression of these genes. Further investigation of Nmus efflux pump genes is required to understand the antibiotic susceptibility pattern of Nmus morphotypes against different classes of antibiotics. This may help us study the effect of antimicrobial resistance upon neisserial persistence at extragenital sites.

*In vivo*, following oral inoculations, 100% of A/J and B6 mice inoculated with the smooth morphotype were persistently colonized. Furthermore, oral burdens of the smooth morphotype were higher in both mouse strains compared to the rough. According to our *in vitro* observations, more pilin is produced by the smooth morphotype than the rough. Additionally, the smooth biofilms show high expression of genes associated with attachment (Table 1). These *in vitro* phenotypes correlate well with our *in vivo* observation that the smooth morphotype can effectively colonize the oral cavity of mice. Another possibility is that Nmus morphotypes may form similar biofilms *in vivo* as observed *in vitro*. In such a case, the rigid and sticky biofilm formed by the smooth morphotype may promote robust colonization of mice. In contrast, due to the fragile nature of the rough biofilms, they may be easily sloughed off and fail to persist as well in the viscous oral environment.

The oral colonization burdens of Nmus morphotypes varied between the two mouse strains used in this study. Both smooth and rough morphotypes showed higher colonization burdens in the A/J mice relative to B6 mice. Higher susceptibility of A/Js compared to B6s has been described in earlier Nmus studies and for other microbial pathogens (15, 16, 46, 47). Thus, variations in Nmus colonization burden may be linked to differences in the host genetics and/or the oral microbiota. Nmus may interact with certain unknown receptors highly or exclusively expressed in the A/Js. Furthermore, colonization by Nmus is known to elicit an antibody response and is affected by host immune responses (16). It is possible that the magnitude or nature of immune response may vary in the two mouse strains used in this study. A comprehensive study comparing transcriptome of the host tissues following Nmus colonization may provide valuable insights about specific genes expressed in each mouse strain and identify host factors promoting higher colonization in the A/J background. Although, the colonization burden is lower in the B6 strain, the smooth morphotype still colonizes 100% of the inoculated B6s. Hence, by using different B6 genetic knockout strains, studies can be performed to determine specific host genetic and immune factors that may affect Nmus colonization.

We observed that the smooth morphotype colonize the guts of more mice as compared to the rough. However, the frequency of gut colonization in A/Js and B6s in our study contrasts with an early observation in CAST mice (15). Differences in the host genetic factors may have affected the gut colonization in these mouse strains. Gut colonization is complex and depends on successful interactions with a heterogenous gut ecosystem. Hence, we can’t exclude the possibility that gut microbiota differences between mouse strains influenced Nmus abundance. Future metagenomics studies may explain the diversity in bacterial populations in this niche and identify organisms that provide some level of colonization resistance to Nmus.

Colonization in the nasal cavity by the Nmus morphotypes was transient and not as robust as observed in the oral cavities. We hypothesize that these differences may be due to variations in the anatomy, physiology, and immune responses in the two microenvironments. Secondly, the composition of local microbiota in these niches may also influence the nasal colonization of Nmus morphotypes (48). Future studies using immunofluorescence microscopy of oral and nasal tissues may elucidate more details about Nmus oral and nasal niches and identify specific host cells colonized by Nmus morphotypes. Following nasal inoculations, both the Nmus morphotypes were able to disseminate to the oral cavities and guts of inoculated mice. Interestingly, within three days of nasal inoculation, the oral burdens of the smooth morphotype were similar to the oral burdens following oral inoculations. Although the exact mechanism of dissemination and factors involved are unknown, our model will provide opportunities to study the *in vivo* dynamics of neisserial dissemination, an important aspect of *Neisseria*-host interactions. However, we cannot rule out the possibility that oral colonization resulted from nasal dripping of the inoculum or self-grooming behavior In conclusion, our study shows that the two Nmus morphotypes show variable phenotypes *in vitro* including biofilm formation, gene expression, vancomycin susceptibility, as well as *in vivo* differences in colonization and persistence. The dynamics of colonization, persistence, and dissemination of Nmus are different depending on the host background, anatomical niches, and inoculation routes. Hence, different combinations of Nmus-host systems may be explored to understand asymptomatic neisserial adaptation and interaction with the host in different niches. The Nmus-mouse system can help identify host and bacterial factors that are essential for stable colonization and persistence in extragenital niches, including natural Nmus orthologs of proteins from human neisserial pathogens, or Nmus recombinants expressing the human pathogen determinants. In the future, this model also holds promise for testing new antimicrobials and vaccine candidates for efficacy against neisserial carriage in the upper respiratory tract.

## MATERIALS AND METHODS

### Strains and primers

All strains and primers used in this study are listed in Table 2.

**TABLE 2.**
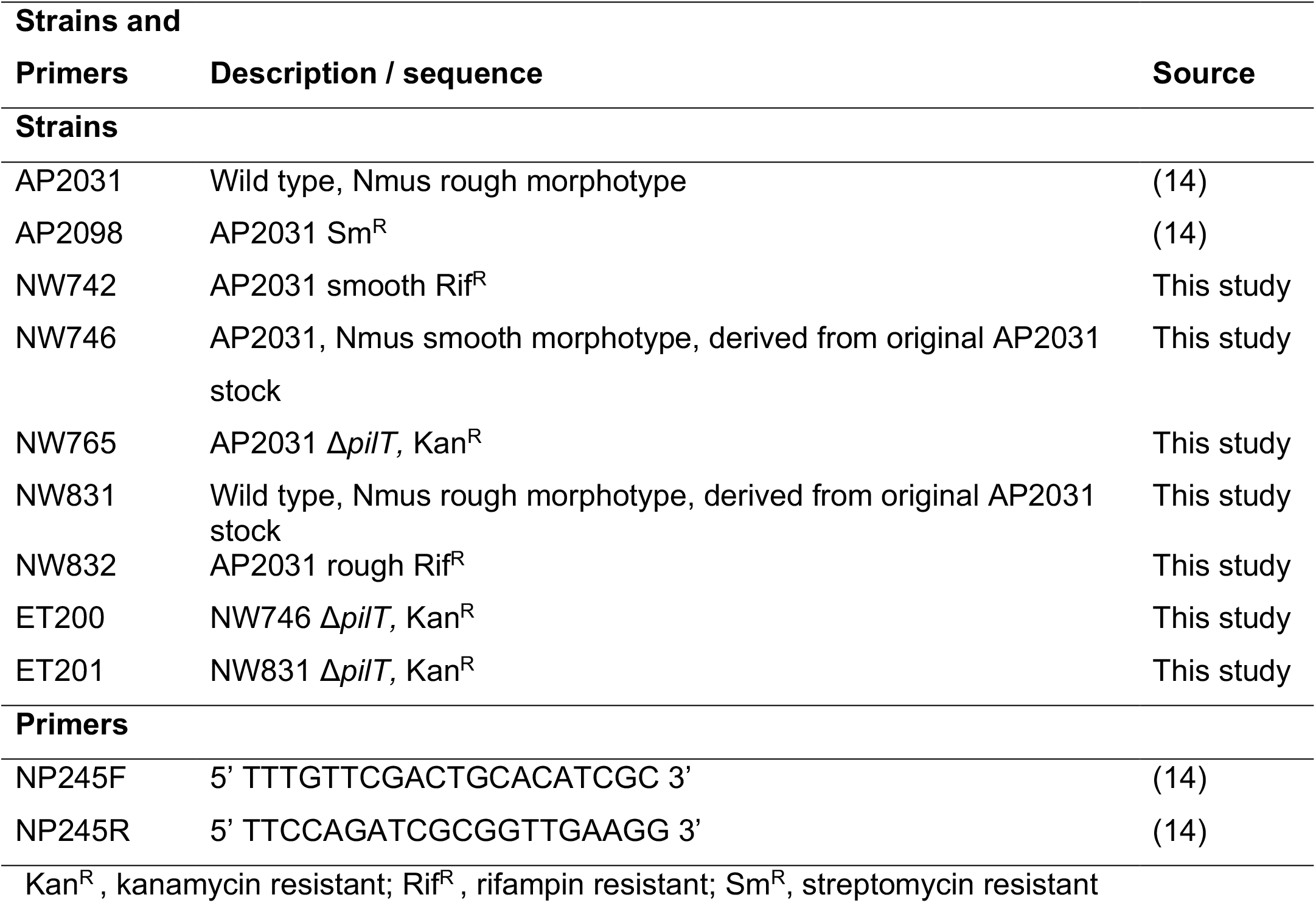
Strains and primers.

### Generation of Nmus strains

NW742 and NW832 are naturally occurring Rif^R^ smooth and rough morphotypes of Nmus type strain, isolated by plating AP2031 on GC medium base (GCB) agar plates containing rifampin (50 mg l^-1^). NW765 is a *pilT* mutant of AP2031 in which *pilT* open reading frame is replaced by a kanamycin resistant cassette (Δ*pilT*::*kan*) as described previously (15). For the generation of ET200 and ET201, primers NP245F and NP245R were used to amplify Δ*pilT*::*kan* locus from NW765. Amplicons were purified and introduced in the smooth and rough morphotypes by spot transformation (49). Transformants were selected on GCB agar plates containing 50 mg l^-1^ kanamycin.

### Bacterial growth conditions

Nmus morphotypes were struck on GCB agar plates with Kellogg’s supplements I and II (50) and incubated at 37°C with 5% CO_2_ for 48 h prior to lawning. When appropriate, rifampin, kanamycin and streptomycin were added in the media at concentrations of 40 mg l^-1^, 50 mg l^-1^ and 100 mg l^-1^ respectively.

### Biofilm formation and fragility assays

Measurement of biofilm formation and fragility was performed as previously described (17) with some modifications. For details, see text in supplemental methods.

### RNA isolation, purification, and transcriptome analysis

Total RNA was extracted from biofilms of Nmus morphotypes using RNeasy Mini kit (Qiagen) using manufacturer’s instructions with some modifications. rRNA depleted RNA samples were used for RNA-Seq library construction and sequencing. For details about library construction, sequencing, and transcriptome analysis, see text in supplemental methods.

### Synteny and sequence analyses

For Nmus and Nme 8013 orthology, shared genes were determined using PanOCT (21) with default suggested parameters with the strict option set to “No”. Gene annotations in Fig. 3 were retrieved from accession numbers FM999788.1, AEQD01000072 (for *macA* and *macB)*, and AAF41681(for *zapE)*. Shared synteny of the *macAB* locus was further confirmed by comparison to Ngo strain FA19 genome annotations (accession number CP012026.1). Protein sequence alignments were performed with Clustal Omega using the default parameters (51). A maximum-likelihood tree of full length pilin sequences was generated based on the Kimura 2-parameter model in MEGA6 (52).

### Pilin extraction and quantification

Extraction of pilin was done by shearing and ultracentrifugation as described previously (25). Pilin preparations were separated by 15% SDS PAGE and stained with Coomassie blue. Image processing and quantification of pilin band intensity was done in ImageJ. For details, see text in supplemental methods.

### Transformation assays

Transformation frequencies of Nmus morphotypes were determined as previously described (14). Genomic DNA from AP2098, a naturally occurring streptomycin resistant strain of Nmus was used as donor DNA (14). For details, see text in supplemental methods.

### Aggregation assay

Aggregation was measured by a sedimentation assay as previously described (53). Bacterial aggregation was indicated as a decrease in absorbance from the initial absorbance. For details, see text in supplemental methods.

### Minimum inhibitory concentration (MIC) measurements

The vancomycin MICs were determined by the Etest method as described previously (54). For details, see text in supplemental methods.

### Mouse inoculations

A/J (stock# 000646) and C57BL/6J (stock# 000664) inbred mice used in the study were obtained from The Jackson Laboratory (Bar harbor, ME). All animal protocols were approved by the Ohio University Institutional Animal Care and Use Committee prior conducting the experiments. For details about mice inoculations and sample processing, see text in supplemental methods.

## ACKNOWLEDGEMENTS

We acknowledge the support and expertise provided by the Ohio University Genomics Facility in completion of the RNA library preparation and MiSeq sequencing. We thank K. Rhodes at the University of Arizona for her suggestions regarding biofilm assays, P. Briaud and R. Carroll at Ohio University for their help with RNA-Seq analyses. We express gratitude for financial support from the Infectious and Tropical Disease Institute, the Molecular and Cellular Biology Graduate Program, and the Department of Biological Sciences, Ohio University. This work was funded in part by NIH (AI124665) and the University of Arizona (College of Medicine) grants awarded to M. So.

## SUPPLEMENTAL MATERIALS

**FIG S1.**
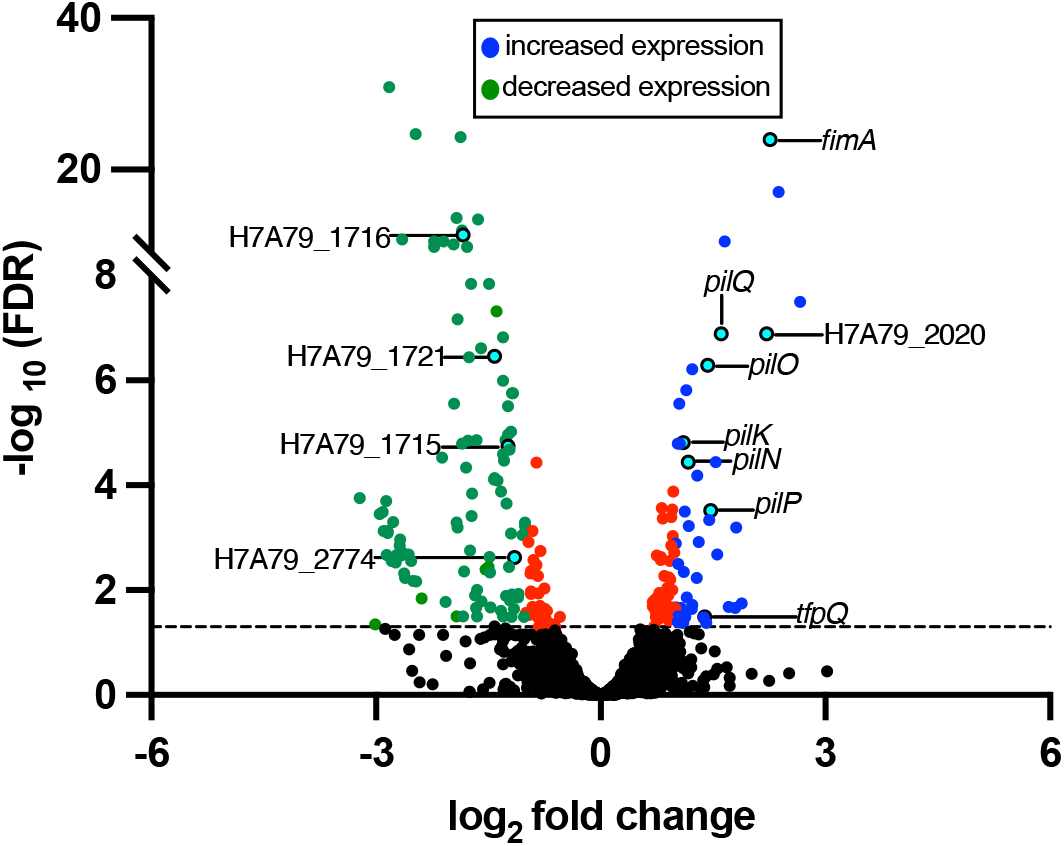
Volcano plot of differentially expressed genes in Nmus biofilms using three sample replicates. Dotted line indicates a False Discovery Rate (FDR) threshold of 0.05. Blue and green dots are genes with significant FDR and log_2_ fold changes for increased (> = 1) and decreased (< = -1) expression in smooth biofilms, respectively. Black dots and red dots represent genes that do not reach the FDR and log_2_ fold change thresholds, respectively. Teal dots with annotations on the right indicate genes that encode factors associated with attachment. Teal dots with annotations on the left denote locus tags for genes that encode efflux pumps.

**FIG S2.**
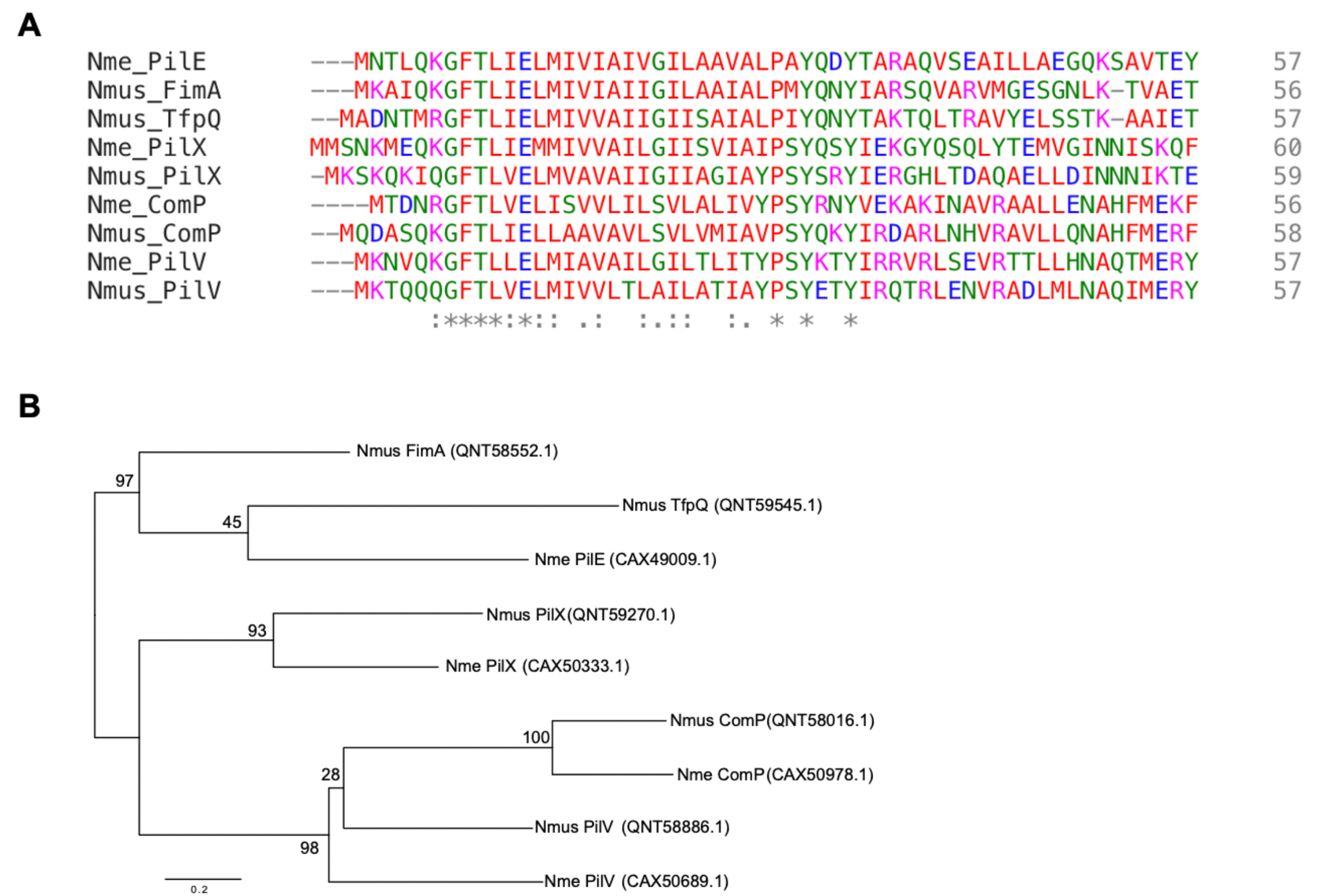
Comparison of pilin sequences from Nme and Nmus. (A) Protein sequence alignments of N-terminal residues of the predicted major and minor pilins in Nme and Nmus. (*), identical residues; (:), conserved residues; (.), semi-conserved residues. (B) Phylogenetic tree of putative pilins. A maximum likelihood tree was constructed using the full-length protein sequences of the putative major and minor pilins in Nme and Nmus. Bootstrap values are indicated (1000 replications). Numbers in the parentheses indicate protein accession numbers. Nme, *Neisseria meningitidis* strain 8013; Nmus, *Neisseria musculi* strain 831. Bar, 0.2 substitutions per site.

**FIG S3.**
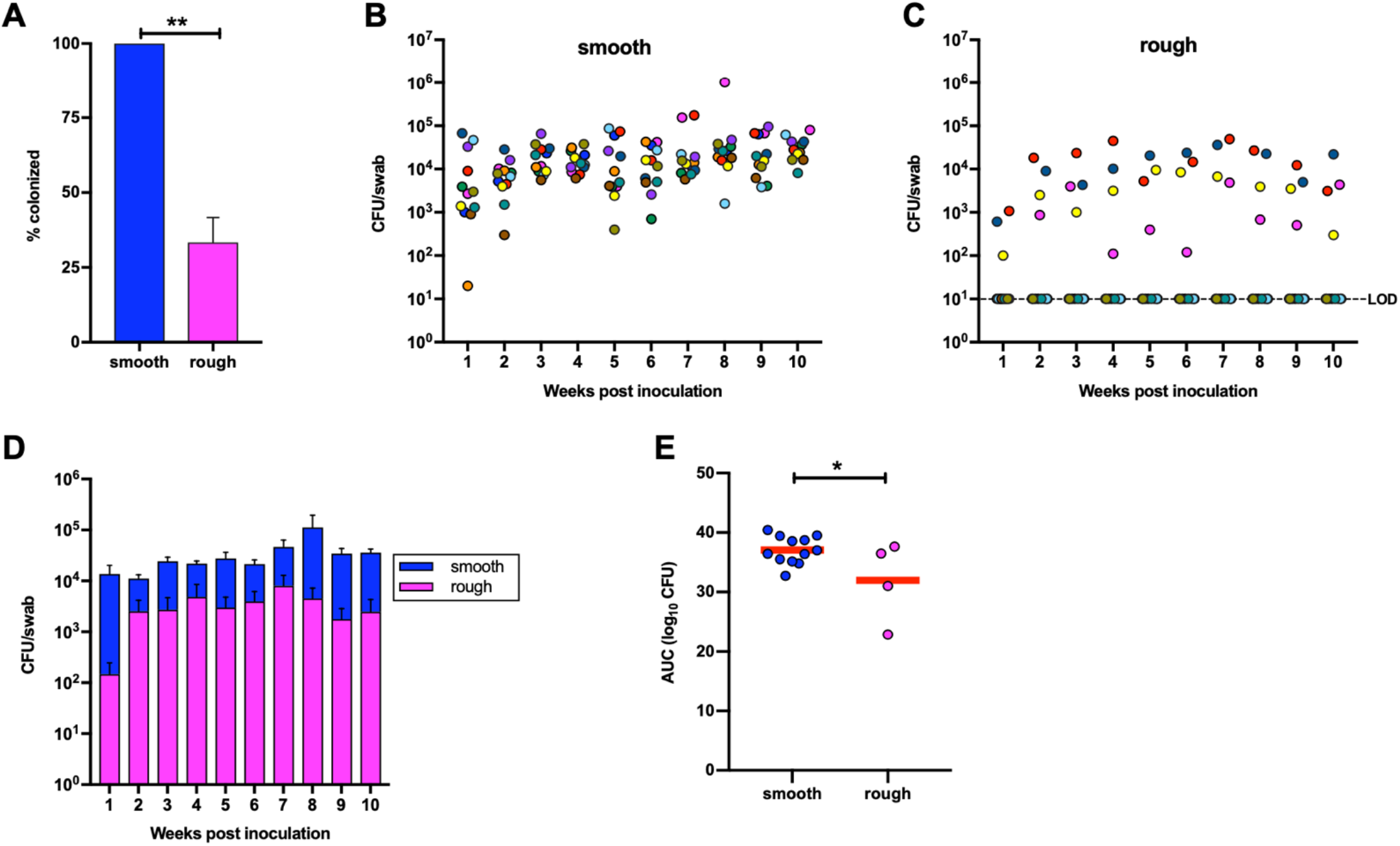
Nmus oral colonization phenotypes in B6 mice. (A) Percentage of B6 mice (n= 12 for both) colonized by Nmus morphotypes following oral inoculations. (B, C) Oral burdens of Nmus morphotypes following oral inoculations. Each colored dot represents a single mouse. LOD, limit of detection. (D) Comparison of the oral burdens of Nmus morphotypes. (E) Cumulative CFU as area under the curve (AUC). The mean AUC (log_10_ CFU versus time) was calculated for each mouse to estimate the bacterial burden over time. The means under the curves were compared between the two morphotypes. Error bars indicate SEM. Red lines indicate means. Statistical analyses for A and D were performed using the Student’s *t* test. *, *P* < 0.05; **, *P* < 0.01.

**FIG S4.**
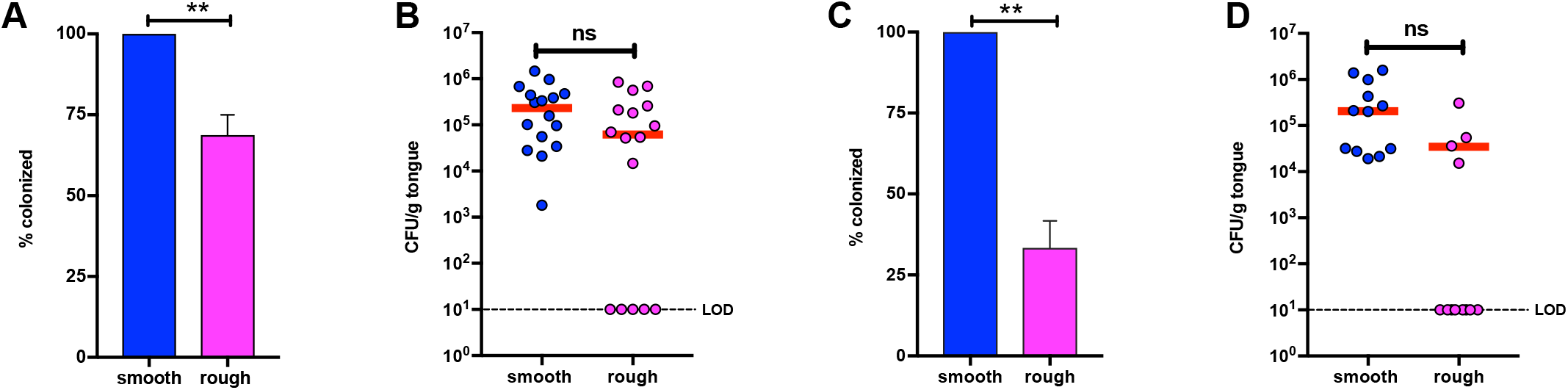
Nmus tongue colonization phenotypes in A/J and B6 mice. (A, C) Percentages of tongues colonized by Nmus morphotypes in (A) A/Js and (C) B6s. Data represent mean with SEM from four (for A/J) and three (for B6) independent experiments. Statistical analysis was performed using the Student’s *t* test. **, *P* < 0.01. (B, D) Tongue CFU burdens for Nmus morphotypes in A/Js (B) and B6s (D). Each dot represents bacterial counts obtained from a tongue harvested from a single mouse. Red lines indicate means. Statistical analysis was performed using the Student’s *t* test with average colonization burden for colonized mice. LOD, limit of detection; ns, *P* > 0.05.

**FIG S5.**
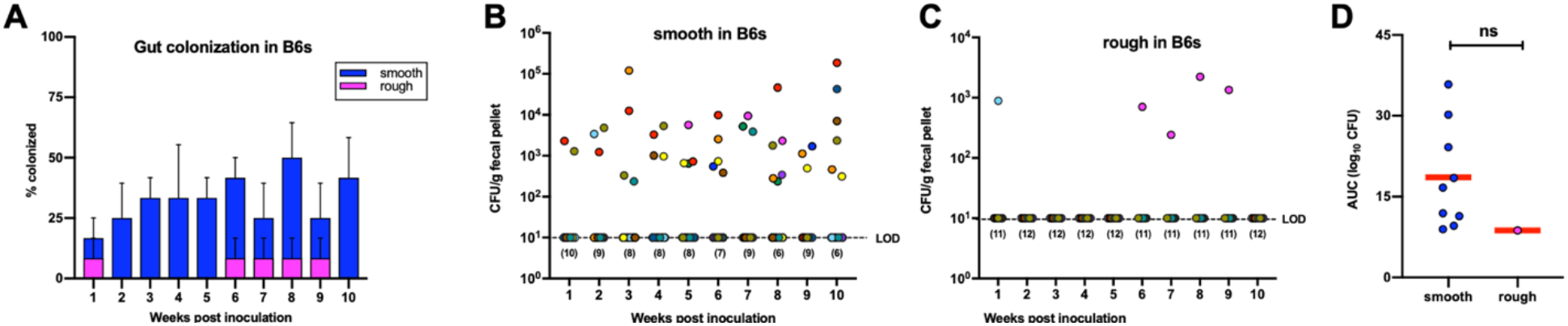
Gut colonization of Nmus morphotypes in B6 mice following oral inoculations. (A) Percentage of B6s colonized by Nmus morphotypes in the gut. Data represent mean with SEM from three independent experiments. Fecal burdens of smooth (B) and rough (C) morphotypes in B6s (n= 12 for both). Each colored dot represents a single mouse. Numbers in parentheses represent the number of mice in which colonization was not detected. LOD, limit of detection. (D) Cumulative CFU as area under the curve (AUC). The mean AUC (log_10_ CFU versus time) was calculated for each mouse to estimate bacterial burden over time. Red lines indicate means. Statistical analysis for D was performed using the Student’s *t* test. ns, *P* > 0.05.

**FIG S6.**
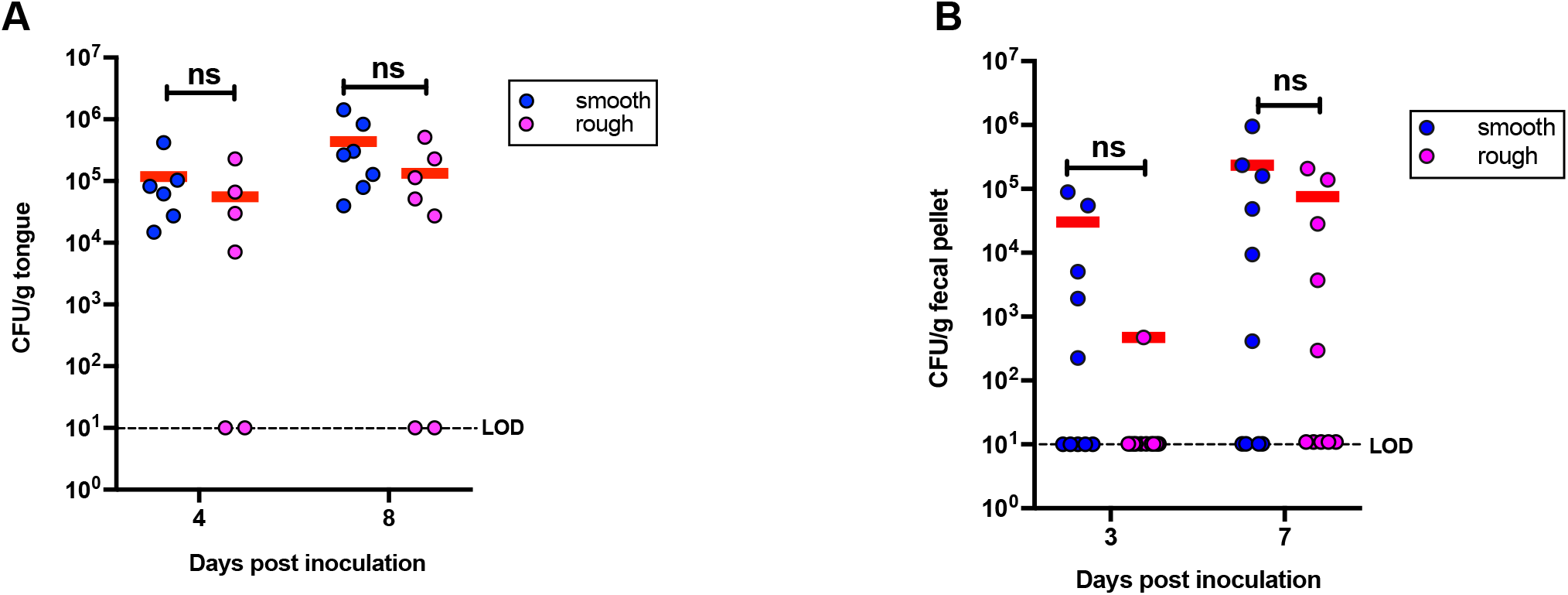
Nmus tongue and gut colonization phenotypes in A/J mice following nasal inoculations. Tongue (A) and gut (B) burdens following nasal inoculations in A/J mice. Each dot represents bacterial counts from a single mouse. Red lines indicate means. LOD, limit of detection. Statistical analyses were performed using the Student’s *t* test on log transformed data. ns, *P* > 0.05.

**Table S1.**
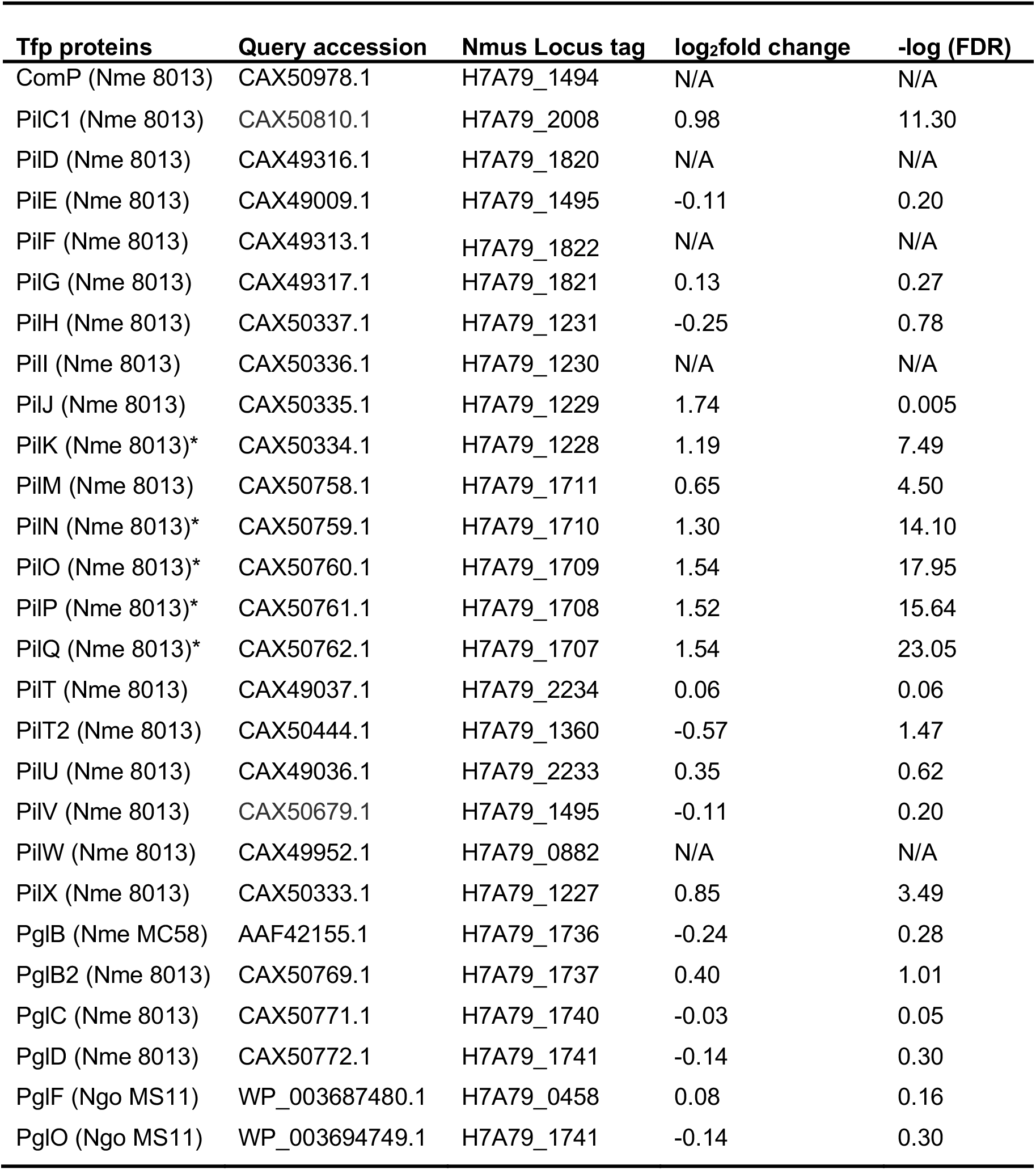
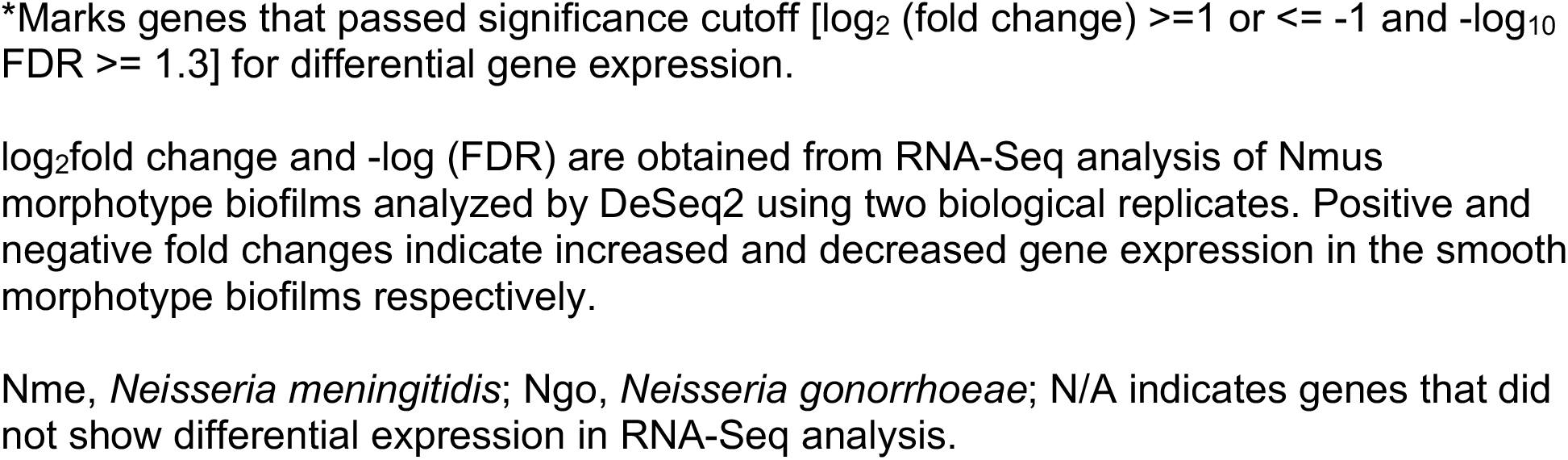
Orthologs of Type IV pilus biogenesis and modification proteins in Nmus

| **Tfp proteins** | **Query accession** | **Nmus Locus tag** | **log<sub>2</sub>fold change** | **-log (FDR)** |
| --- | --- | --- | --- | --- |
| ComP (Nme 8013) | CAX50978.1 | H7A79_1494 | N/A | N/A |
| PilC1 (Nme 8013) | CAX50810.1 | H7A79_2008 | 0.98 | 11.30 |
| PilD (Nme 8013) | CAX49316.1 | H7A79_1820 | N/A | N/A |
| PilE (Nme 8013) | CAX49009.1 | H7A79_1495 | -0.11 | 0.20 |
| PilF (Nme 8013) | CAX49313.1 | H7A79_1822 | N/A | N/A |
| PilG (Nme 8013) | CAX49317.1 | H7A79_1821 | 0.13 | 0.27 |
| PilH (Nme 8013) | CAX50337.1 | H7A79_1231 | -0.25 | 0.78 |
| PilI (Nme 8013) | CAX50336.1 | H7A79_1230 | N/A | N/A |
| PilJ (Nme 8013) | CAX50335.1 | H7A79_1229 | 1.74 | 0.005 |
| PilK (Nme 8013)* | CAX50334.1 | H7A79_1228 | 1.19 | 7.49 |
| PilM (Nme 8013) | CAX50758.1 | H7A79_1711 | 0.65 | 4.50 |
| PilN (Nme 8013)* | CAX50759.1 | H7A79_1710 | 1.30 | 14.10 |
| PilO (Nme 8013)* | CAX50760.1 | H7A79_1709 | 1.54 | 17.95 |
| PilP (Nme 8013)* | CAX50761.1 | H7A79_1708 | 1.52 | 15.64 |
| PilQ (Nme 8013)* | CAX50762.1 | H7A79_1707 | 1.54 | 23.05 |
| PilT (Nme 8013) | CAX49037.1 | H7A79_2234 | 0.06 | 0.06 |
| PilT2 (Nme 8013) | CAX50444.1 | H7A79_1360 | -0.57 | 1.47 |
| PilU (Nme 8013) | CAX49036.1 | H7A79_2233 | 0.35 | 0.62 |
| PilV (Nme 8013) | CAX50679.1 | H7A79_1495 | -0.11 | 0.20 |
| PilW (Nme 8013) | CAX49952.1 | H7A79_0882 | N/A | N/A |
| PilX (Nme 8013) | CAX50333.1 | H7A79_1227 | 0.85 | 3.49 |
| PglB (Nme MC58) | AAF42155.1 | H7A79_1736 | -0.24 | 0.28 |
| PglB2 (Nme 8013) | CAX50769.1 | H7A79_1737 | 0.40 | 1.01 |
| PglC (Nme 8013) | CAX50771.1 | H7A79_1740 | -0.03 | 0.05 |
| PglD (Nme 8013) | CAX50772.1 | H7A79_1741 | -0.14 | 0.30 |
| PglF (Ngo MS11) | WP_003687480.1 | H7A79_0458 | 0.08 | 0.16 |
| PglO (Ngo MS11) | WP_003694749.1 | H7A79_1741 | -0.14 | 0.30 |

\*Marks genes that passed significance cutoff [ $\log_2$ (fold change) $\geq 1$ or $\leq -1$ and $-\log_{10}$ FDR $\geq 1.3$ ] for differential gene expression.
$\log_2$ fold change and $-\log$ (FDR) are obtained from RNA-Seq analysis of Nmus morphotype biofilms analyzed by DeSeq2 using two biological replicates. Positive and negative fold changes indicate increased and decreased gene expression in the smooth morphotype biofilms respectively.
Nme, *Neisseria meningitidis*; Ngo, *Neisseria gonorrhoeae*; N/A indicates genes that did not show differential expression in RNA-Seq analysis.

**Table S3.**
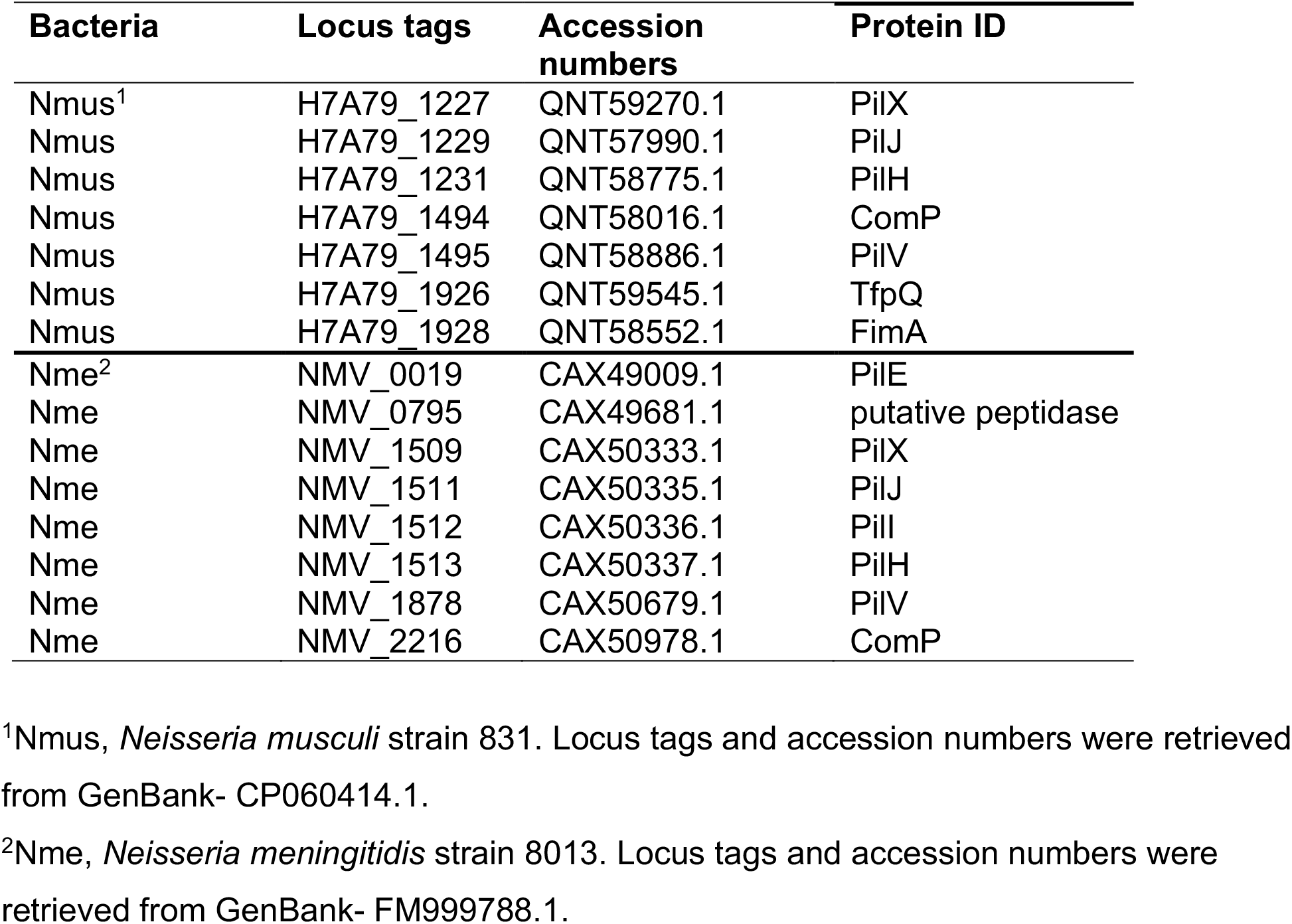
Putative pilins identified by PilFind program in Nmus and Nme

## SUPPLEMENTAL METHODS

### Biofilm formation and fragility assays

Eighteen hours old lawns of Nmus morphotypes were suspended in GC broth and adjusted to an optical density (OD_600_) of 0.05. Five hundred microliters of the suspension was added to each well of a 24 well flat bottom polystyrene plate (CytoOne) and incubated statically at 37°C with 5% CO_2_ for 16 h. Following incubation, the medium was removed, and biofilms were air dried briefly until no wet spots were visible. Dried biofilms were stained with 0.1% (w/v) crystal violet for 10 min followed by washing three times gently with 1 ml of phosphate-buffered saline (PBS) to remove non-adherent crystal violet. Retained Crystal Violet was solubilized in 1 ml of 30% glacial acetic acid and the OD_595_ was measured. Experiments were performed three times with technical triplicates.

Measurement of biofilm fragility was performed similarly except that the biofilms were not air dried before staining with Crystal Violet. Experiments were performed three times with technical quadruplicates.

### RNA isolation, purification, and transcriptome analysis

Total RNA was extracted from biofilms of Nmus morphotypes. Biofilms were grown in 90 mm petri dishes for 16 h and washed twice with PBS to remove planktonic bacteria. One ml of RNAlater™ Stabilization Solution (Thermofisher) was added to each plate and biofilms were scraped and collected in sterile microfuge tubes. The tubes were centrifuged for 3 min at 14, 000 x g at 4 °C to pellet the cells. Each pellet was suspended in 100 ml Tris EDTA buffer pH 8.0 containing 1 mg ml^-1^ of lysozyme and incubated for 5 min at room temperature with vortexing every 1 min. RNA extraction was completed using RNeasy Mini kit (Qiagen) using the manufacturer’s instructions. DNA was removed by DNase digestion using the Turbo DNA-free kit (Invitrogen). RNA concentrations and integrity was measured using an Agilent 2100 bioanalyzer (Agilent Technologies). Only RNAs with RIN values greater than 9.5 were used for further processing. Ribosomal RNA was removed by the Ribo-Zero rRNA Removal Kit for bacteria (Illumina) per manufacturer’s instructions. RNA quality was checked again in the bioanalyzer to ensure removal of rRNA. Three biofilm RNA samples were prepared from each morphotype for analysis.

RNA-Seq library construction and sequencing of rRNA depleted RNA samples was performed at the Ohio University Genomic Facility. Libraries were constructed using Illumina Truseq Stranded mRNA Library Prep kit (Illumina Cat. # 20020594) per manufacturer’s instruction starting with the Fragment, Prime, and Finish step to accommodate non-polyadenylated, rRNA-depleted mRNA. Sequencing was performed with Illumina Miseq reagent kit v3, 150 cycle kit (Illumina, Cat. #MS-102-3001) on the Illumina MiSeq system to obtain 75 pair-end reads according to the manufacturer’s guide. RNA-Seq samples were demultiplexed and exported. Illumina reads were mapped using Bowtie (1) to the genome of Nmus strain NW831 (2). Gene expression counts for all samples were estimated using HTseq (3). Counts were then tabularized and imported into R studio for analyses. Differentially expressed genes were analyzed using the DESeq2 R package (4) and filtered using an FDR cutoff of < = 0.05 and an absolute log_2_ fold change of >=1, for the comparison of the two Nmus morphotypes. Principal Component Analyses (PCAs) were generated using various R packages based on normalized Variance Stabilized Transformation counts from DESeq2. Differentially expressed genes were plotted on a Volcano plot using GraphPad Prism 9. RNA-Seq data were deposited in the Gene Expression Omnibus database under the accession number GSE184230.

### Pilin extraction and quantification

Lawns of Nmus strains grown on GCB agar plates for 18 h were resuspended in 50 mM CHES (2-(Cyclohexylamino) ethanesulfonic acid) pH 9.5. Bacterial suspensions were vortexed for 2 min and cells were pelleted at 18,000 × g for 10 min. The supernatant was collected and spun down at 100,000 × g for 2 h in Optima TLX ultracentrifuge (Beckman Coulter). Pelleted pilin was resuspended in 50 mM CHES. Preparations were separated by 15% SDS PAGE and stained with Coomassie blue. Image processing and quantification of pilin band intensity was done with ImageJ. The density of pilin bands from the smooth morphotype were set as 1 to measure the relative pilin densities from other strains. Adjusted density was calculated by dividing the relative density of pilin bands from each strain to the relative density of an approximately 60 kDa shared band as a loading control. Experiments were performed three times.

### Transformation assays

Suspensions of Nmus morphotypes were prepared in GC broth with 5 mM MgSO_4_ and adjusted to OD_600_ of 1.5. Thirty microliters of this suspension were added to 0.2 ml GC broth with 5 mM MgSO_4_ and 1 μg of donor DNA. The mixture was incubated for 15 min at 37°C and added to 2 ml GC broth containing supplements I and II for further incubation at 37°C with 5% CO_2_ for 3 h. Following incubation, transformation mixture was plated on GCB agar plates to count the total CFU and GCB agar plates with streptomycin (100 mg l^-1^) to count streptomycin resistant (Sm^R^) transformants. Transformation frequency was calculated as the ratio of Sm^R^ CFU to total CFU per μg of DNA. Transformation mixtures without added donor DNA served as negative controls. Experiments were repeated three times.

### Aggregation assay

Nmus morphotypes grown on GCB agar plates for 18 h were suspended in PBS and adjusted to OD_600_ of 1.0. Suspensions were vortexed for 10 sec and allowed to stand statically. After each 15 min, 100 μl of each suspension was taken from the top of the tube and the OD_600_ was measured until 90 min. Bacterial aggregation was indicated as a decrease in absorbance from the initial absorbance. Experiments were repeated three times.

### Minimum inhibitory concentration (MIC) measurements

Nmus morphotypes were harvested from GCB agar plates, suspended in GC broth, and diluted to OD_600_ of 1.0. Bacterial suspensions were applied with sterile polystyrene swabs (Fisher Scientific) to GCB agar. Plates were allowed to dry briefly, and vancomycin Etest strips (Liofilchem) were placed on the agar surface. The plates were then incubated at 37°C in 5% CO_2_ for 18 h. MICs were interpreted as the point where the elliptical edge intersected the strip. MICs were measured at least three times for each isolate.

### Mouse inoculations

Five-week-old female A/J or B6 mice were acclimatized in the Ohio University animal facility for two weeks prior to use. One week before the inoculations, the oral cavities of mice were swabbed with BD BBL™ Culture Swab (Becton, Dickinson and Company) and plated to detect presence of any *Neisseria* spp. No *Neisseria* spp. were detected in any mice prior to our experiments.

For oral inoculations, rifampin resistant strains of Nmus morphotypes were harvested from GCB agar plates after 18 h, suspended in PBS, and adjusted to an OD_600_ of 1.0. Mice were restrained manually and 50 μl of bacterial suspension was slowly inoculated into the oral cavity of each mouse. Oral swabs were collected weekly for 10 weeks. Swabs were suspended in GCB + 20% glycerol, vortexed for 1 min, diluted in GC broth and plated on GCB agar plates containing rifampin. Plates were incubated at 37°C with 5% CO_2_ for 48 h prior to colony enumeration.

Fecal pellets were collected in sterile microfuge tubes and weighed. One ml of GCB + 20% glycerol was added to each pellet, followed by suspension with a sterile plain wooden applicator. Pellet suspensions were vortexed for 1 min and plated to enumerate CFU per gram of fecal pellet.

For nasal inoculations, bacterial inoculum was prepared similarly but with an OD_600_ of 1.5. Mice were restrained manually and 20 μl of bacterial suspension (10 μl per nare) was slowly inoculated into the nasal cavities of mice. Oral swabs were collected and processed from the nasally inoculated mice after 3 days post inoculation (dpi) or 7 dpi as described above. At 4 dpi or 8 dpi, mice were euthanized by CO_2_ asphyxiation. The nasal tip was snipped with sterile scissors and the nasal cavity was swabbed with a saline pre-wetted Puritan™ calcium alginate swab. Nasal swabs were suspended in 1 ml GCB + 20% glycerol, processed, and plated as for oral swabs described above.

To harvest the tongue, incisions were made on either side of the mouth of euthanized mice. The oral cavity was opened, and the tongue was held with sterile forceps from the tip and cut at the base with sterile scissors. Each tongue was cut into small pieces, homogenized in a Mini Beadbeater-16 (BioSpec Products), diluted in GCB, and plated to enumerate CFU per gram of tissue.

## REFERENCES

1. Humbert MV, Christodoulides M. 2019. Atypical, Yet Not Infrequent, Infections with Neisseria Species. Pathogens 9.

2. Liu G, Tang CM, Exley RM. 2015. Non-pathogenic Neisseria: members of an abundant, multi-habitat, diverse genus. Microbiology (Reading) 161:1297–1312.

3. Donati C, Zolfo M, Albanese D, Tin Truong D, Asnicar F, Iebba V, Cavalieri D, Jousson O, De Filippo C, Huttenhower C, Segata N. 2016. Uncovering oral Neisseria tropism and persistence using metagenomic sequencing. Nat Microbiol 1:16070.

4. Diallo K, Trotter C, Timbine Y, Tamboura B, Sow SO, Issaka B, Dano ID, Collard J-M, Dieng M, Diallo A, Mihret A, Ali OA, Aseffa A, Quaye SL, Bugri A, Osei I, Gamougam K, Mbainadji L, Daugla DM, Gadzama G, Sambo ZB, Omotara BA, Bennett JS, Rebbetts LS, Watkins ER, Nascimento M, Woukeu A, Manigart O, Borrow R, Stuart JM, Greenwood BM, Maiden MCJ. 2016. Pharyngeal carriage of Neisseria species in the African meningitis belt. J Infect 72:667–677.

5. Rotman E, Seifert HS. 2014. The Genetics of Neisseria Species. Ann Rev Genet 48:405–431.

6. Moxon ER, Jansen VAA. 2005. Phage variation: understanding the behaviour of an accidental pathogen. Trends Microbiol 13:563–565.

7. Barbee LA. 2014. Preparing for an era of untreatable gonorrhea. Curr Opin Infect Dis 27:282–287.

8. Callaghan MM, Dillard JP. 2019. Mucus is a key factor in Neisseria meningitidiscommensalism. mSphere 4.

9. Dorey RB, Theodosiou AA, Read RC, Jones CE. 2019. The nonpathogenic commensal Neisseria: friends and foes in infectious disease. Curr Opin Infect Dis 32:490–496.

10. Deasy AM, Guccione E, Dale AP, Andrews N, Evans CM, Bennett JS, Bratcher HB, Maiden MC, Gorringe AR, Read RC. 2015. Nasal Inoculation of the commensal Neisseria lactamica Inhibits carriage of Neisseria meningitidis by young adults: A Controlled Human Infection Study. Clin Infect Dis 60:1512–20.

11. Custodio R, Johnson E, Liu G, Tang CM, Exley RM. 2020. Commensal Neisseria cinerea impairs Neisseria meningitidis microcolony development and reduces pathogen colonisation of epithelial cells. PLoS Pathog 16:e1008372.

12. Kim WJ, Higashi D, Goytia M, Rendón MA, Pilligua-Lucas M, Bronnimann M, McLean JA, Duncan J, Trees D, Jerse AE, So M. 2019. Commensal Neisseria kill Neisseria gonorrhoeae through a DNA-Dependent Mechanism. Cell Host Microbe 26:228–239.e8.

13. Marri PR, Paniscus M, Weyand NJ, Rendón MA, Calton CM, Hernández DR, Higashi DL, Sodergren E, Weinstock GM, Rounsley SD, So M. 2010. Genome sequencing reveals widespread virulence gene exchange among human Neisseria species. PLoS One 5:e11835.

14. Weyand NJ, Ma M, Phifer-Rixey M, Taku NA, Rendón MA, Hockenberry AM, Kim WJ, Agellon AB, Biais N, Suzuki TA, Goodyer-Sait L, Harrison OB, Bratcher HB, Nachman MW, Maiden MCJ, So M. 2016. Isolation and characterization of Neisseria musculi sp. nov., from the wild house mouse. Int J Syst Evol Microbiol 66:3585–3593.

15. Ma M, Powell DA, Weyand NJ, Rhodes KA, Rendón MA, Frelinger JA, So M. 2018. A natural mouse model for Neisseria colonization. Infect Immun 86:e00839–17.

16. Powell DA, Ma M, So M, Frelinger JA. 2018. The commensal Neisseria musculi modulates host innate immunity to promote oral colonization. ImmunoHorizons 2:305.

17. Merritt JH, Kadouri DE, O’Toole GA. 2005. Growing and analyzing static biofilms. Curr Protoc Microbiol Chapter 1:Unit 1B.1.

18. Kolappan S, Coureuil M, Yu X, Nassif X, Egelman EH, Craig L. 2016. Structure of the Neisseria meningitidis Type IV pilus. Nat Commun 7:13015.

19. Duan Q, Nandre R, Zhou M, Zhu G. 2017. Type I fimbriae mediate in vitro adherence of porcine F18ac+ enterotoxigenic Escherichia coli (ETEC). Ann Microbiol 67:793–799.

20. Punsalang AP, Jr., Sawyer WD. 1973. Role of pili in the virulence of Neisseria gonorrhoeae. Infect Immun 8:255–63.

21. Fouts DE, Brinkac L, Beck E, Inman J, Sutton G. 2012. PanOCT: automated clustering of orthologs using conserved gene neighborhood for pan-genomic analysis of bacterial strains and closely related species. Nucleic Acids Res 40:e172.

22. Rusniok C, Vallenet D, Floquet S, Ewles H, Mouzé-Soulama C, Brown D, Lajus A, Buchrieser C, Médigue C, Glaser P, Pelicic V. 2009. NeMeSys: a biological resource for narrowing the gap between sequence and function in the human pathogen Neisseria meningitidis. Genome Biol 10:R110.

23. Carbonnelle E, Hélaine S, Prouvensier L, Nassif X, Pelicic V. 2005. Type IV pilus biogenesis in Neisseria meningitidis: PilW is involved in a step occurring after pilus assembly, essential for fibre stability and function. Mol Microbiol 55:54–64.

24. Imam S, Chen Z, Roos DS, Pohlschröder M. 2011. Identification of surprisingly diverse type IV pili, across a broad range of gram-positive bacteria. PLoS One 6:e28919.

25. Higashi DL, Biais N, Weyand NJ, Agellon A, Sisko JL, Brown LM, So M. 2011. N. elongata produces type IV pili that mediate interspecies gene transfer with N. gonorrhoeae. PLoS One 6:e21373.

26. Dietrich M, Mollenkopf H, So M, Friedrich A. 2009. Pilin regulation in the pilT mutant of Neisseria gonorrhoeae strain MS11. FEMS Microbiol Lett 296:248–56.

27. Aas FE, Wolfgang M, Frye S, Dunham S, Løvold C, Koomey M. 2002. Competence for natural transformation in Neisseria gonorrhoeae: components of DNA binding and uptake linked to type IV pilus expression. Mol Microbiol 46:749–760.

28. Long CD, Tobiason DM, Lazio MP, Kline KA, Seifert HS. 2003. Low-level pilin expression allows for substantial DNA transformation competence in Neisseria gonorrhoeae. Infect Immun 71:6279–91.

29. Hélaine S, Carbonnelle E, Prouvensier L, Beretti JL, Nassif X, Pelicic V. 2005. PilX, a pilus-associated protein essential for bacterial aggregation, is a key to pilus-facilitated attachment of Neisseria meningitidis to human cells. Mol Microbiol 55:65–77.

30. Thapa E, Aluvathingal J, Nadendla S, Mehta A, Tettelin H, Weyand NJ. 2021. Complete genome sequence of Neisseria musculi using Illumina and PacBio Sequencing. Microbiol Resour Announc 10:e0045221.

31. Blight MA, Pimenta AL, Lazzaroni JC, Dando C, Kotelevets L, Séror SJ, Holland IB. 1994. Identification and preliminary characterization of temperature-sensitive mutations affecting HlyB, the translocator required for the secretion of haemolysin (HlyA) from Escherichia coli. Mol Gen Genet 245:431–40.

32. Reyes Ruiz LM, Williams CL, Tamayo R. 2020. Enhancing bacterial survival through phenotypic heterogeneity. PLoS Pathog 16:e1008439.

33. Sousa A, Machado I, Pereira MO. Phenotypic switching: an opportunity to bacteria thrive, p. In (ed),

34. Weiser JN. 1993. Relationship between colony morphology and the life cycle of Haemophilus influenzae: the contribution of lipopolysaccharide phase variation to pathogenesis. J Infect Dis 168:672–80.

35. Diaz L, Hoare A, Soto C, Bugueno I, Silva N, Dutzan N, Venegas D, Salinas D, Perez-Donoso JM, Gamonal J, Bravo D. 2015. Changes in lipopolysaccharide profile of Porphyromonas gingivalis clinical isolates correlate with changes in colony morphology and polymyxin B resistance. Anaerobe 33:25–32.

36. Zelewska MA, Pulijala M, Spencer-Smith R, Mahmood HA, Norman B, Churchward CP, Calder A, Snyder LAS. 2016. Phase variable DNA repeats in Neisseria gonorrhoeae influence transcription, translation, and protein sequence variation. Microb Genom 2:e000078.

37. Wanford JJ, Green LR, Aidley J, Bayliss CD. 2018. Phasome analysis of pathogenic and commensal Neisseria species expands the known repertoire of phase variable genes, and highlights common adaptive strategies. PLoS One 13:e0196675.

38. Brown DR, Helaine S, Carbonnelle E, Pelicic V. 2010. Systematic functional analysis reveals that a set of seven genes is involved in fine-tuning of the multiple functions mediated by type IV pili in Neisseria meningitidis. Infect Immun 78:3053–63.

39. Wolfgang M, van Putten JP, Hayes SF, Dorward D, Koomey M. 2000. Components and dynamics of fiber formation define a ubiquitous biogenesis pathway for bacterial pili. Embo j 19:6408–18.

40. Assalkhou R, Balasingham S, Collins RF, Frye SA, Davidsen T, Benam AV, Bjørås M, Derrick JP, Tønjum T. 2007. The outer membrane secretin PilQ from Neisseria meningitidis binds DNA. Microbiology (Reading) 153:1593–1603.

41. Cheng Y, Johnson MD, Burillo-Kirch C, Mocny JC, Anderson JE, Garrett CK, Redinbo MR, Thomas CE. 2013. Mutation of the conserved calcium-binding motif in Neisseria gonorrhoeae PilC1 impacts adhesion but not piliation. Infect Immun 81:4280–9.

42. Rudel T, Boxberger HJ, Meyer TF. 1995. Pilus biogenesis and epithelial cell adherence of Neisseria gonorrhoeae pilC double knock-out mutants. Mol Microbiol 17:1057–71.

43. Ryll RR, Rudel T, Scheuerpflug I, Barten R, Meyer TF. 1997. PilC of Neisseria meningitidis is involved in class II pilus formation and restores pilus assembly, natural transformation competence and adherence to epithelial cells in PilC-deficient gonococci. Mol Microbiol 23:879–92.

44. Auda IG, Ali Salman IM, Odah JG. 2020. Efflux pumps of Gram-negative bacteria in brief. Gene Reports 20:100666.

45. Schlör S, Schmidt A, Maier E, Benz R, Goebel W, Gentschev I. 1997. In vivo and in vitro studies on interactions between the components of the hemolysin (HlyA) secretion machinery of Escherichia coli. Mol Gen Genet 256:306–19.

46. Domínguez-Punaro Mde L, Segura M, Radzioch D, Rivest S, Gottschalk M. 2008. Comparison of the susceptibilities of C57BL/6 and A/J mouse strains to Streptococcus suis serotype 2 infection. Infect Immun 76:3901–10.

47. Tuite A, Elias M, Picard S, Mullick A, Gros P. 2005. Genetic control of suceptibility to Candida albicans in susceptible A/J and resistant C57BL/6J mice. Genes & Immunity 6:672–682.

48. Hardy BL, Merrell DS. 2021. Friend or Foe: Interbacterial Competition in the Nasal Cavity. J Bacteriol 203.

49. Dillard JP. 2011. Genetic Manipulation of Neisseria gonorrhoeae. Current protocols in microbiology Chapter 4:Unit4A.2-Unit4A.2.

50. Kellogg DS, Jr., Peacock WL, Jr., Deacon WE, Brown L, Pirkle DI. 1963. Neisseria gonorrhoeae I. Virulence genetically linked to clonal variation. J Bact 85:1274–1279.

51. Sievers F, Wilm A, Dineen D, Gibson TJ, Karplus K, Li W, Lopez R, McWilliam H, Remmert M, Söding J, Thompson JD, Higgins DG. 2011. Fast, scalable generation of high-quality protein multiple sequence alignments using Clustal Omega. Mol Syst Biol 7:539–539.

52. Tamura K, Stecher G, Peterson D, Filipski A, Kumar S. 2013. MEGA6: Molecular Evolutionary Genetics Analysis version 6.0. Mol Biol Evol 30:2725–2729.

53. Trunk T, Khalil HS, Leo JC. 2018. Bacterial autoaggregation. AIMS microbiology 4:140–164.

54. Thapa E, Knauss HM, Colvin BA, Fischer BA, Weyand NJ. 2020. Persistence dynamics of antimicrobial resistant Neisseria in the pharynx of rhesus macaques. Antimicrob Agents Chemother 64: e02232–19.

## REFERENCES

1. Langmead B, Trapnell C, Pop M, Salzberg SL. 2009. Ultrafast and memory-efficient alignment of short DNA sequences to the human genome. Genome Biol 10:R25.

2. Thapa E, Aluvathingal J, Nadendla S, Mehta A, Tettelin H, Weyand NJ. 2021. Complete Genome Sequence of Neisseria musculi Using Illumina and PacBio Sequencing. Microbiol Resour Announc 10:e0045221.

3. Anders S, Pyl PT, Huber W. 2015. HTSeq--a Python framework to work with high-throughput sequencing data. Bioinformatics 31:166–9.

4. Love MI, Huber W, Anders S. 2014. Moderated estimation of fold change and dispersion for RNA-seq data with DESeq2. Genome Biol 15:550.

